# Genome-wide dysregulation of histone acetylation in the Parkinson’s disease brain

**DOI:** 10.1101/785550

**Authors:** Lilah Toker, Gia T Tran, Janani Sundaresan, Ole-Bjørn Tysnes, Guido Alves, Kristoffer Haugarvoll, Gonzalo S Nido, Christian Dölle, Charalampos Tzoulis

## Abstract

Parkinson disease (PD) is a complex neurodegenerative disorder of largely unknown etiology. While several genetic risk factors have been identified, the involvement of epigenetics in the pathophysiology of PD is mostly unaccounted for. We conducted a histone acetylome-wide association study in PD, using brain tissue from two independent cohorts of cases and controls. Immunoblotting revealed increased acetylation at several histone sites in PD, with the most prominent change observed for H3K27, a marker of active promoters and enhancers. Chromatin immunoprecipitation sequencing (ChIP-seq) further indicated that H3K27 hyperacetylation in the PD brain is a genome-wide phenomenon, with a strong predilection for genes implicated in the disease, including *SNCA, PARK7, PRKN* and *MAPT*. Integration of the ChIP-seq with transcriptomic data revealed that the correlation between promoter H3K27 acetylation and gene expression is attenuated in PD patients, suggesting that H3K27 acetylation may be decoupled from transcription in the PD brain. Our findings strongly suggest that dysregulation of histone acetylation plays an important role in the pathophysiology of PD and identify novel epigenetic signatures associated with the disease.

## Introduction

Parkinson’s disease (PD) is the second most common neurodegenerative disorder, affecting ∼1.8 % of the population above the age of 65 years^1^. The neuropathological hallmark of PD is the progressive loss of dopaminergic neurons in the *substantia nigra pars compacta* (SNc) in the presence of α-synuclein-positive inclusions termed Lewy pathology. Additional neurodegenerative changes occur in multiple regions of the nervous system, including several brainstem nuclei, the olfactory bulb, hippocampus, amygdala and the neocortex, as well as the autonomic and enteric nervous systems^2^.

The etiology of PD remains largely unknown. While monogenic forms of PD exist, they are rare, generally accounting for less than 5-10% of all cases in most populations. The vast majority of patients have “idiopathic PD”, which is of complex etiology^3^. Genome-wide association studies (GWAS) have identified multiple risk *loci* associated with idiopathic PD. These make a relatively small collective contribution to the disease risk, however, and have not been linked to specific molecular mechanisms that could be targeted therapeutically^4^.

Epigenetic modifications, such as DNA methylation and histone acetylation, modulate gene expression without altering the underlying DNA sequence. Dysregulation of histone acetylation has been previously linked to aging^5^ and neurodegeneration^6^ and changes in histone acetylation at specific genomic regions have been reported in Alzheimer’s disease^7,8^. However, despite the emerging role of epigenetics in disease initiation and progression, only a few small-scale studies of histone acetylation in PD have been reported to date, providing inconclusive results^9–11^. Moreover, the genomic landscape of histone acetylation in PD has never been assessed.

Here, we report the first genome-wide study of histone acetylation in the PD brain, based on two independent cohorts from the Norwegian ParkWest study (PW)^12^ and the Netherlands Brain Bank (NBB). Our findings reveal that H3K27 acetylation is profoundly dysregulated at a genome-wide level in the PD brain.

## Results

### Increased histone acetylation and histone protein levels in the PD brain

We assessed global protein acetylation in the prefrontal cortex (PFC) of 13 individuals with PD and 13 demographically matched controls from the Park West (PW) cohort by immunoblot analysis using an antibody against acetylated lysine (Fig. 1A, Supplementary Table S1). We concentrated on PFC, because while affected by the disease, this region is less confounded by altered cell composition due to neurodegeneration and neuroinflammation^2^. Linear regression analysis of the densitometric measurements, adjusted for age, sex and PMI indicated an increased acetylation signal at ∼17 kDa, consistent with the molecular weight of core histone proteins (p = 0.014, Fig. 1A).

**Fig. 1:**
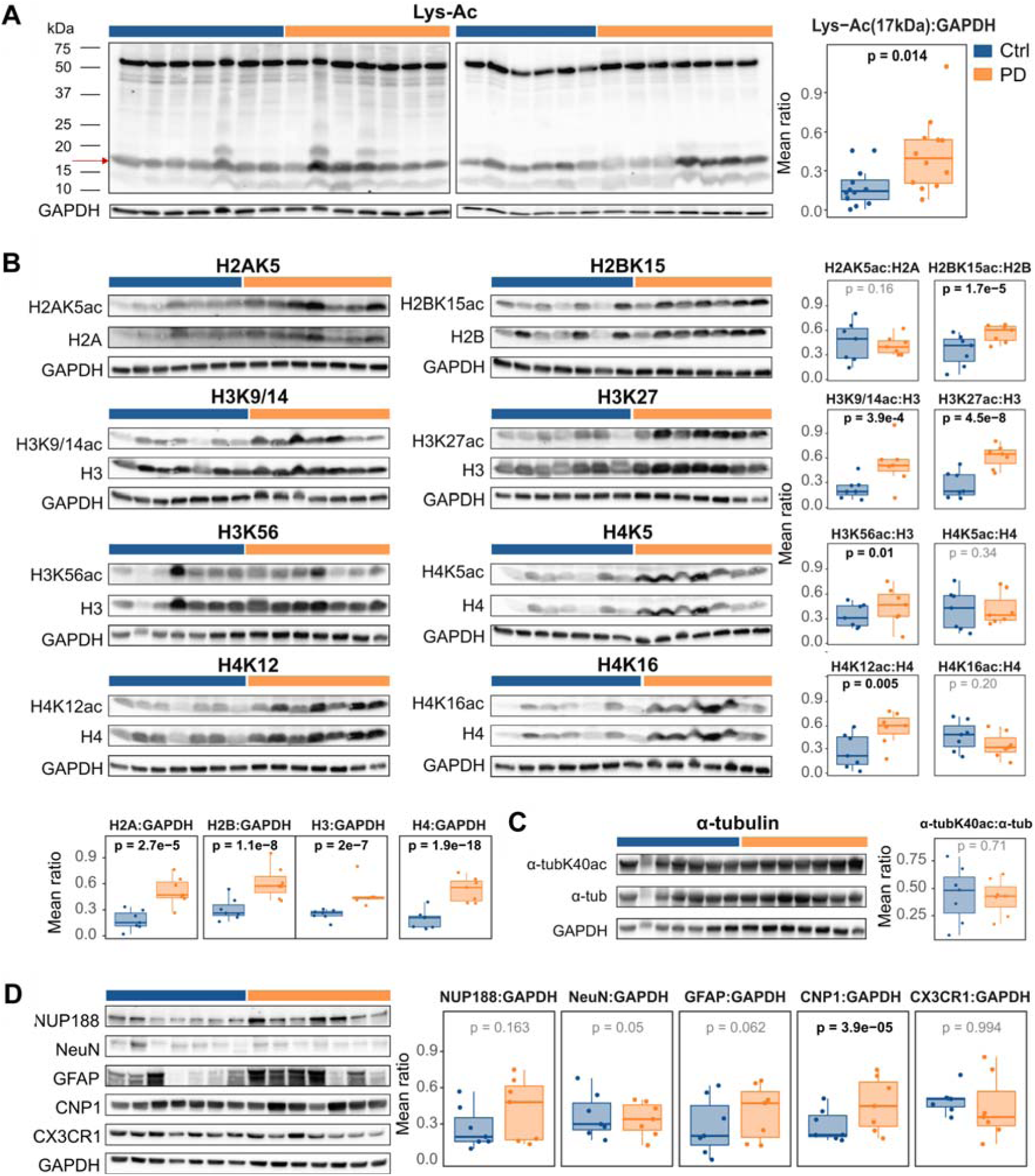
Hyperacetylation of multiple histone sites in the PD prefrontal cortex. Representative Western blots and the ratios of their quantified densitometric measurements of the indicated proteins (or their modifications) in individuals with PD (orange) and controls (blue) are shown. GAPDH served as loading control. Values were rescaled [0,1]. Boxplots show the mean of replicates for each individual. The indicated p-values are based on a linear mixed model (see methods). **A.** Global protein acetylation detected by a pan-acetyl lysine antibody. **B.** Relative acetylation of histone lysine residues and quantification of the total histone proteins. **C.** Relative acetylation of α-tubulin on lysine K40. **D**, Immunodetection and densitometric quantification of relative protein levels of the nuclear marker (NUP188) and cell type-specific marker proteins: NeuN – neurons, GFAP – astrocytes, CNP1 – oligodendrocytes, CX3CR1 - microglia.

To further dissect histone- and site-specificity, we analyzed the acetylation state at specific sites on the four core histones: H2AK5ac, H2BK15ac, H3K9/K14ac, H3K27ac, H3K56ac, H4K5ac, H4K12ac, and H4K16ac. The acetylation status of α-tubulin, a cytoskeletal protein typically regulated by acetylation, was assessed as a non-histone control. To maximize sensitivity, we focused our analyses on the 7 PD and 7 control samples showing the strongest difference in the pan-acetylation signal (Supplementary Table S1). The acetylated fraction of each histone site as well as total histone quantity were analyzed using linear mixed models with gender, age and PMI as fixed effects and subject and blot number as random effects. These analyses indicated a significant increase in H2BK15ac (p = 1.4 x 10^−5^), H3K9/14ac (p = 3.9 x 10^−4^), H3K27ac (p = 4.5 x 10^−8^), H3K56ac (p = 0.01) and H4K12ac (p = 5.0 x 10^−3^) in PD. No significant difference was found for H2AK5, H4K5, H4K16, or α-tubulin (Fig. 1B-C, Supplementary Table S2). In addition, we found increase in the total quantity of the four histone proteins (H2A: p = 2.7 x 10^−5^, H2B: p = 1.1 x 10^−8^, H3: p = 2.0 x 10^−7^; H4: p = 3.9 x 10^−18^).

To access the cellular composition of our samples, we carried out immunoblot analyses of markers for neurons (NeuN), astrocytes (GFAP), oligodendrocytes (CNP1), microglia (CX3CR1) and total number of nuclei (NUP188), in the same samples used for acetylation assessment. We found a significant increase in CNP1 (p = 3.9 x 10^−5^, Fig. 1D, Supplementary Table S2), but no changes in the other markers. To ensure that our findings were not driven by differences in oligodendrocyte abundance, we repeated the analyses adding CNP1 as a covariate to the statistical model. As shown in Supplementary Table S2, all of the indicated hyperacetylated sites remained significant after CNP1 adjustment.

To determine the regional specificity of altered H3K27ac and H3K9/14ac in the PD brain, we assessed these two markers in the cerebellum and striatum from the same patients. H3K27 was significantly hyperacetylated in the PD striatum (p = 0.034) and cerebellum (p = 0.016). H3K9/14 was significantly hypoacetylated in the PD cerebellum (p = 7.0 x 10^−3^) and unchanged in the striatum (p = 0.7, Fig. 2, Supplementary Table S2).

**Fig. 2:**
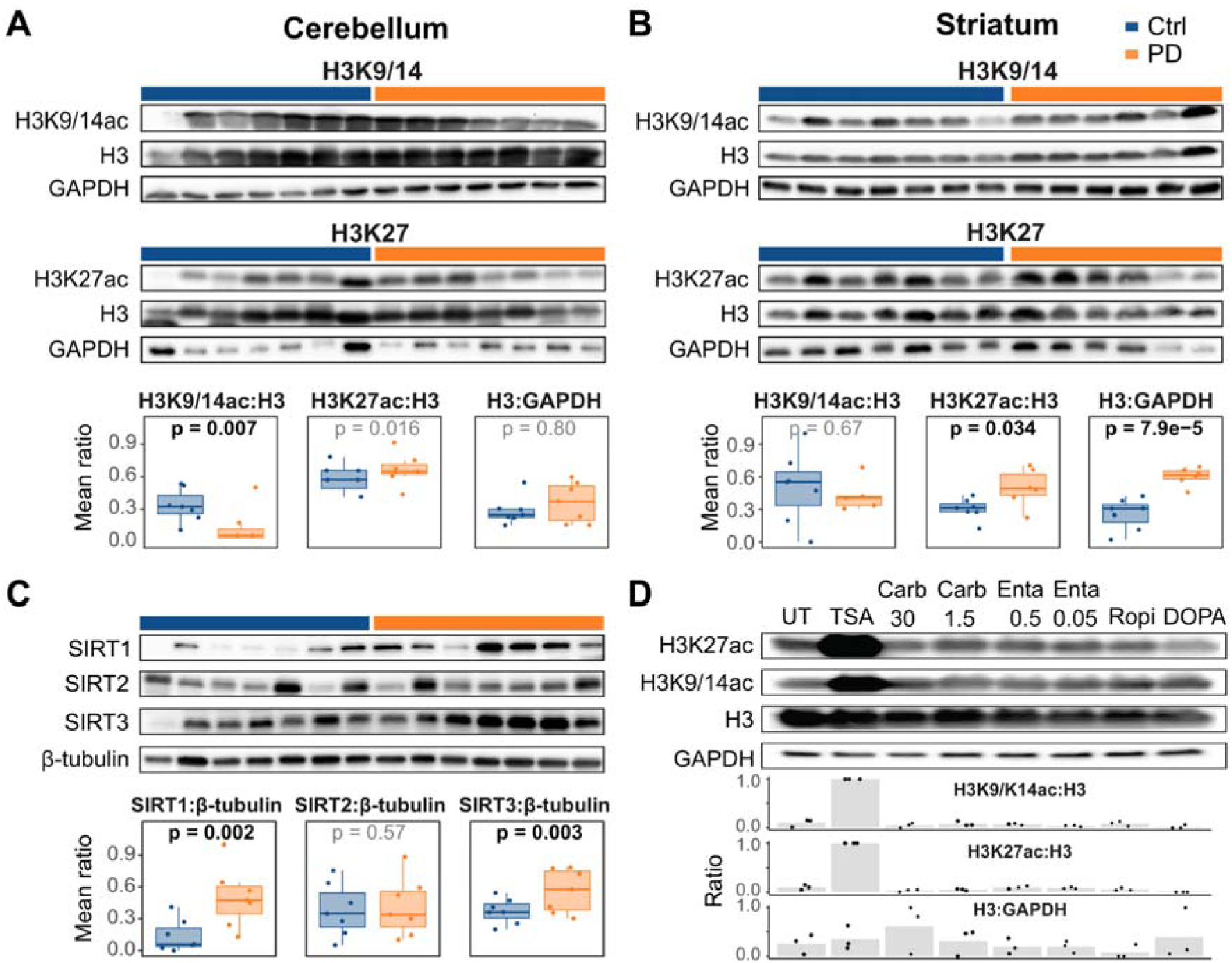
Altered H3 acetylation occurs in multiple brain regions, is accompanied by sirtuin upregulation and is not induced by anti-Parkinson drugs in-vitro. **A-C**, Representative Western blots and quantification plots showing the ratios of quantified densitometric measurements of the indicated proteins in individuals with PD (orange) and controls (blue). GAPDH and β-tubulin serve as loading controls. Values were rescaled [0,1]. Boxplots show the mean of replicates for each individual. The indicated p-values are based on a linear mixed model (see methods). **A, B**, Acetylated H3K9/14 and H3K27 and total histone protein H3 levels in the cerebellum (A) and striatum (B) from individuals with PD and controls. **C.** Sirtuin proteins in the prefrontal cortex of PD and controls. **D**, Acetylated H3K9/14, H3K27 and total H3 protein in differentiated SH-SY5Y-derived dopaminergic neuron-like cells treated with the histone deacetylase inhibitor Trichostatin A (TSA) or anti-Parkinson drugs used by the PW patients during their last year of life. The numbers indicate the concentration (in µM) of the drug. Carb: Carbidopa; Enta: Entacapone; Ropi: Ropinirole; DOPA: L-Dopa. UT: untreated control. A representative blot is shown. Bar plots show the mean quantified ratios of three biological replicates (rescaled [0,1]). Dots indicate the values of each of the individual replicates.

### Histone hyperacetylation in the PD brain is accompanied by upregulation of sirtuin protein levels

The PD brain is characterized by widespread mitochondrial complex-I deficiency^13,14^. It has been shown that complex-I deficiency can result in histone hyperacetylation^10,15,16^ due to a decreased NAD^+^/NADH ratio attenuating the activity of NAD^+^-dependent class III histone deacetylases known as sirtuins^15,17–20^. We thus postulated that the observed histone hyperacetylation may be mediated by reduced sirtuin activity. Direct confirmation of this hypothesis would require a reliable determination of intracellular NAD^+^, which was not feasible in our samples due to rapid postmortem degradation^21^. However, it was previously reported that decreased sirtuin activity caused by NAD^+^ deficiency is accompanied by increase in the protein quantity^22,23^. Thus, we assessed the protein levels of SIRT1, SIRT2 and SIRT3, localized to the nucleus, cytosol and mitochondria, respectively, as surrogates for their activity. We were specifically interested in these proteins because they have been linked to aging and neurodegeneration^17^. Immunoblot analyses indicated a significant upregulation of SIRT1 (p = 2.0 x 10^−3^) and SIRT3 (p = 3.0 x 10^−3^) in PD samples. No difference was detected for SIRT2 (Fig. 2C, Supplementary Table S2), in line with the unaltered acetylation state of its target, α-tubulin (Fig. 1C).

### Histone hyperacetylation is not induced by anti-Parkinson drugs *in cellulo*

We next investigated whether the observed altered histone acetylation in PD could result from treatment with anti-Parkinson agents. To test this, we exposed fully differentiated SH-SY5Y cells to the drugs used by most individuals with PD from the PW cohort during their last year of life (L-DOPA, carbidopa, entacapone and ropinirole), and monitored acetylation of H3K9/14 and H3K27. Immunoblot analysis revealed no effect for any of the tested anti-Parkinson drugs on histone acetylation (Fig. 2D).

### Acetylated H3K27 exhibits higher genomic occupancy in PD

H3K27ac exhibited the most pronounced alteration in our samples. To gain insight into the genomic distribution of this modification, we carried out ChIP-seq analysis of this modification in PFC samples from the PW cohort (Supplementary Table S1). First, we characterized the genomic distribution of H3K27ac in the PD and control groups. For this purpose, we identified H3K27ac enriched genomic regions (peaks) in each group separately (see Methods). In line with the increased H3K27ac signal observed in the immunoblot analysis, a higher number of peaks (146,763 vs. 135,662), as well as a higher proportion of group-unique peaks (32.2% vs. 13.2%), and wider genome coverage (10.6% vs. 9.11%) were found in the PD-compared to the control group (Fig. 3A, Supplementary Table S3). We then compared the distribution of genomic annotations and detection p-values of overlapping and unique peaks between the PD and control groups. In both groups, unique peaks occurred more frequently in intergenic and intronic regions than in promoter and exonic regions (Supplementary Fig. S1A,B). Since unique peaks were also characterized by significantly higher detection p-values, compared to the overlapping peaks (Fig. S1C,D), we believe that most of these represent random H3K27 acetylation.

**Fig. 3:**
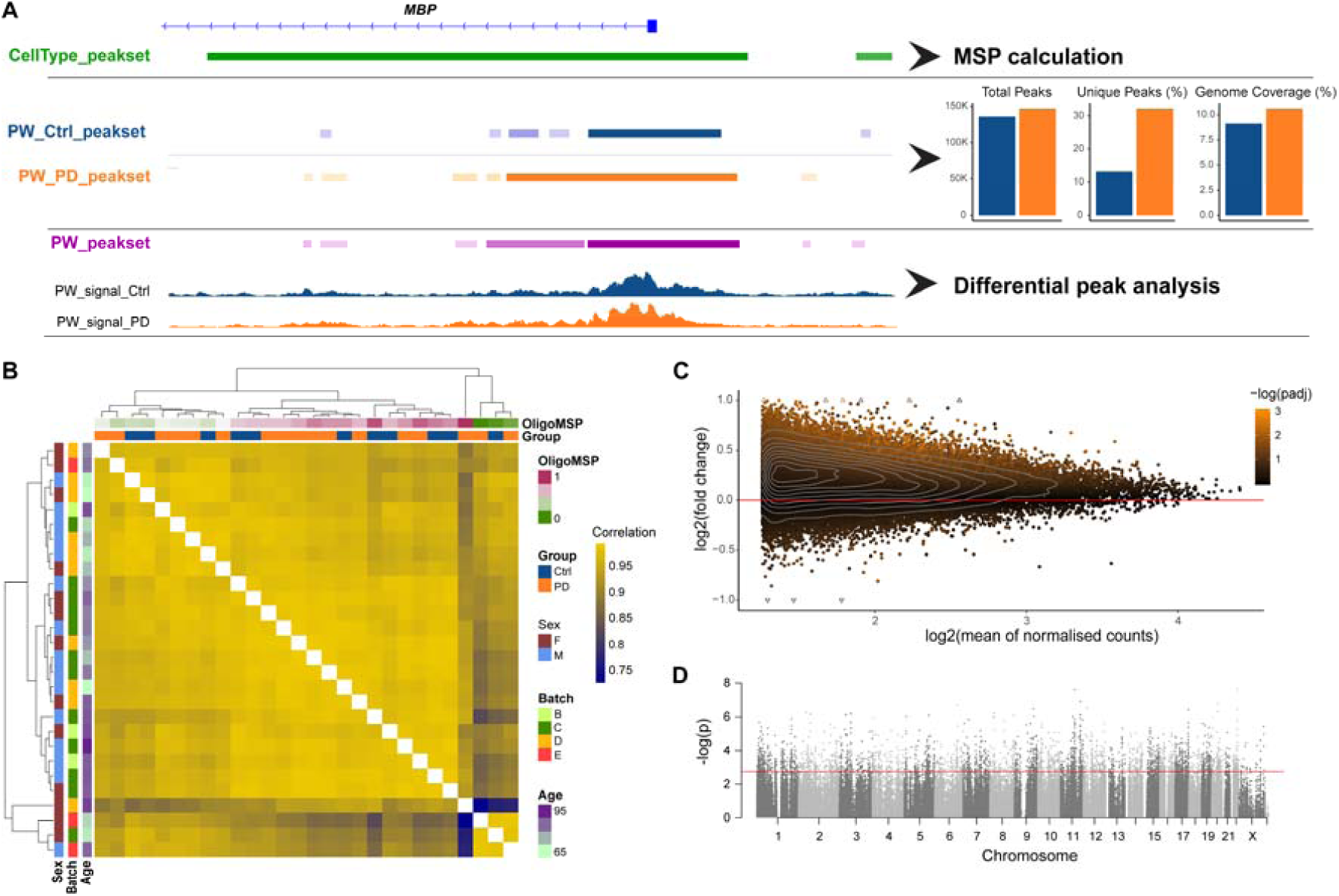
H3K27ac ChIP-seq analysis of PD and control samples. **A.** Different peak-sets used for the different stages of ChIP-seq analysis. Shown, as an example, is a genomic region around the oligodendrocyte marker *MBP*. Peaks annotated to this region, in each peak-set are shown by colored boxes. The CellType_peak-set was used for the MSP calculation. Group specific peak-sets were used to characterize the distribution of H3K27ac bound regions in cases and controls. Barplots show the comparison between the two groups (Supplementary Table S3). PD group exhibited higher number of peaks, higher percentage of unique peaks and higher percentage of genomic coverage, consistent with genome-wide H3K27 hyperacetylation. The PW_peak-set was used for the differential peak analysis. Shown are group overlays of sample fold of change compared to input control. **B.** Hierarchical clustering of the samples indicated that samples cluster based on their cellular composition. Supplementary Fig. S6 shows the association of each of the variables with the main principal components of the data. **C.** MA plot based on the PW_peak-set indicates that increased H3K27ac in PD is observed genome-wide, rather than being restricted to specific regions. **D**, Manhattan plot showing the distribution of genomic locations and differential p-values of the H3K27ac PW_peak-set. The red dashed line indicates the -log10 of the highest p-value with < 5% false positive rate (Benjamini-Hochberg adjusted p-value < 0.05). Corresponding plots for the NBB cohort are shown in Supplementary Fig. S5.

### ChIP-seq data corroborates H3K27 hyperacetylation in the PD brain

Next, we redefined H3K27ac peaks by identifying H3K27ac enriched regions in aggregated samples (see Methods). The final peak-set consisted of 160,521 peaks, covering 11.6% of the genome and distributed over all 22 autosomes and both sex chromosomes (median peak size, 817bp, Fig. 3D). All downstream analyses were performed based on this peak-set, henceforth referred to as PW_peak-set.

ChIP-seq data reflects the abundance and genomic distribution of the target protein (or protein modification) which can vary substantially between samples^24^. Based on the observed increase in H3K27ac fraction in the immunoblot analysis, we anticipated an increase in overall H3K27ac ChIP-seq signal in PD. Thus, we reasoned that the most suitable approach for normalization of our data would be using reads in peaks (RiP) annotated to house-keeping genes^24^. We modified this approach by manually selecting peaks annotated to known housekeeping genes and exhibiting a high and broad peak signal covering most of the gene body. The rationale here was that these genes should be saturated with histone acetylation and are, therefore, expected to exhibit little biological variation. Supplementary Fig. S2 shows the H3K27ac ChIP-seq peaks annotated to the previously proposed^24^ and the chosen housekeeping genes.

To validate our normalization approach, we performed additional H3K27ac immunoblot and ChIP-seq analyses in samples from 55 individuals (including the study subjects, Supplementary Table S1) and tested the correlation between the immunoblot and ChIP-seq data normalized by different approaches. As shown in Supplementary Fig. S3, the highest correlation between immunoblotting and ChIP-seq data was observed when the total number of RiP was normalized to our manually chosen peaks. Importantly, comparison of normalized RiP between PD and control samples corroborated the finding of general H3K27 hyperacetylation in PD (Supplementary Fig. S3).

### Cell composition is a major source of variation in bulk tissue H3K27ac ChIP-seq data

Since different cell-types exhibit distinct epigenetic landscapes^25^, bulk tissue data must be evaluated in the context of the underlying cellular composition^26–28^. To estimate the cellular composition in our ChIP-seq data, we used an approach similar to the one described by Mancarci et al.^27^. Briefly, we intersected genomic H3K27ac binding sites differentiating between NeuN-positive and NeuN-negative brain cells^25^ with the list of brain cell-type specific marker-genes^27^, in order to obtain genomic H3K27ac cell-type marker sites. The first principal component of reads aligned to the marker sites was used to obtain Marker Site Profiles (MSPs) and was used as a proxy for the relative abundance of the corresponding cell-types across the samples. We validated this approach in publicly available ChIP-seq data from cortical and cerebellar bulk tissue samples^29^, and ChIP-seq data from entorhinal cortex of individuals with Alzheimer’s disease and healthy controls^8^. MSP analysis of these data successfully recapitulated the well-known increased abundance of glial cells in cortical vs cerebellar samples^30,31^, and decreased number of neurons in the entorhinal cortex of subjects with Alzheimer’s disease compared to controls^32,33^ (Supplementary Fig. S4A,B).

Next, we performed MSP analysis in the PW_peak-set data. This analysis indicated no significant differences in cell-type abundance between the PD and control groups (Supplementary Fig. S5). Nevertheless, due to the variable oligodendrocyte content in the subset of subjects tested by immunoblotting, we chose to assess the effect of oligodendrocyte MSPs together with other experimental and demographic covariates on the ChIP-seq data. Strikingly, sample-to-sample correlation indicated that the samples clustered mainly based on oligodendrocytes MSPs (Fig. 3B). Accordingly, the first principal component of the data was mostly associated with oligodendrocyte MSP (Supplementary Fig. S6). These results confirmed that oligodendrocyte composition is a major source of variance in our data. Therefore, oligodendrocyte MSPs were accounted for in all downstream analyses.

### Hyperacetylation of H3K27 in the PD brain is a genome-wide phenomenon

We next sought to identify differentially acetylated regions (DARs) for H3K27 between PD and control samples in our discovery cohort. After filtering (see Methods), 133,716 peaks annotated to 17,182 genes remained for downstream analyses. Five samples (one control and four PD samples) were detected as outliers and excluded from differential peak analysis (Supplementary Table S1). Differential peak analysis was performed using the “DESeq2” R package, adjusting for age, sex, batch, PMI, oligodendrocyte MSP and normalization peak ratio. DARs were defined as differentially acetylated peaks with adjusted p-value < 0.05.

Our analysis identified 2,877 hyperacetylated and only 14 hypoacetylated PD-associated DARs, annotated to 1,434 and 9 genes, respectively. The top-ranked hyperacetylated and hypoacetylated DARs resided within the *DLG2* (adjusted p = 9.8 x 10^−4^) and *PTPRH* (adjusted p-value 9.0 x 10^−3^) genes, respectively, both of which have been previously associated with PD^4,34–36^. The full list of DARs with their annotated genes is provided in Supplementary Table S4. An MA plot (the log ratio vs. mean count) of all regions included in the analysis indicated a genome-wide trend for H3K27 hyperacetylation in PD (Fig. 3C).

### H3K27 hyperacetylation in the PD cortex is replicated in an independent cohort

To validate our findings, we carried out ChIP-seq in an independent cohort from the Netherlands Brain Bank (NBB, Supplementary Table S1). In agreement with our finding from the PW cohort, we observed an increase in the percentage of unique peaks as well as increased total genomic coverage in the PD group (Supplementary table S3). The distribution of genomic annotations among the unique and overlapping peaks and the distribution of peak p-values were similar between the two cohorts (Supplementary Fig. S1). As in the PW cohort, cellular composition accounted for most of the interindividual variance in this cohort (Supplementary Fig. S6B, S7A).

Differential acetylation analysis corroborated the finding of PD-associated genome-wide H3K27 hyperacetylation observed in the PW cohort (Supplementary Fig. S5B). In total, the analysis identified 2,486 hyperacetylated DARs, mapped to 946 genes, and 227 hypoacetylated DARs, mapped to 253 genes (Supplementary Table S5). Strikingly, 275 out of the 946 genes with hyperacetylated DARs and 2 out of 253 genes with hypoacetylated DARs replicated across both cohorts (Fig. 4C, hypergeometric test: p = 2.29 x 10^−85^ and p = 7.9 x 10^−^ 3, respectively). These 277 genes are henceforth referred to as “replicated genes”. The top replicated genes (ranked based on their adjusted p-value in the PW cohort) are shown in Table 1 and a complete list is provided in Supplementary Table S6.

**Table 1.** Top replicated genes between PW and NBB cohorts.

| GeneSymbol | log2(FC)_PW | padj_PW | log2(FC)_NBB | padj_NBB |
| --- | --- | --- | --- | --- |
| Hypoacetylated genes |  |  |  |  |
| *PTPRH* | -0.81 | 9.1E-03 | -0.94 | 0.03 |
| *JUP* | -0.43 | 0.05 | -0.44 | 0.02 |
| Hyperacetylated genes |  |  |  |  |
| *DLG2* | 0.68 | 9.8E-04 | 0.77 | 2.0E-03 |
| *TNRC6B* | 0.74 | 9.8E-04 | 0.9 | 0.03 |
| *SIK3* | 0.66 | 3.2E-03 | 0.52 | 0.01 |
| *SGK1* | 0.65 | 3.2E-03 | 0.72 | 0.01 |
| *PPP2R2B* | 1.3 | 3.5E-03 | 0.7 | 8.0E-03 |
| *FCHSD2* | 0.83 | 3.5E-03 | 1 | 0.01 |
| *CTNND2* | 0.52 | 3.5E-03 | 0.63 | 0.01 |
| *PTK2* | 0.71 | 3.5E-03 | 0.55 | 0.02 |
| *SLCO1A2* | 0.58 | 3.5E-03 | 0.68 | 0.02 |
| *LPAR1* | 0.99 | 3.5E-03 | 0.55 | 0.04 |
| *FRMD4B* | 0.64 | 3.5E-03 | 0.53 | 0.04 |
| *RNF216* | 0.6 | 5.1E-03 | 0.59 | 2.3E-03 |
| *TMEM132B* | 0.88 | 5.1E-03 | 0.59 | 0.02 |
| *PIP4K2A* | 0.57 | 5.1E-03 | 0.52 | 0.03 |
| *MAP7* | 0.55 | 5.9E-03 | 0.52 | 0.04 |
| *PDE4B* | 0.59 | 6.5E-03 | 0.67 | 2.4E-03 |
| *FRMD5* | 0.55 | 6.5E-03 | 0.67 | 0.01 |

**Fig. 4:**
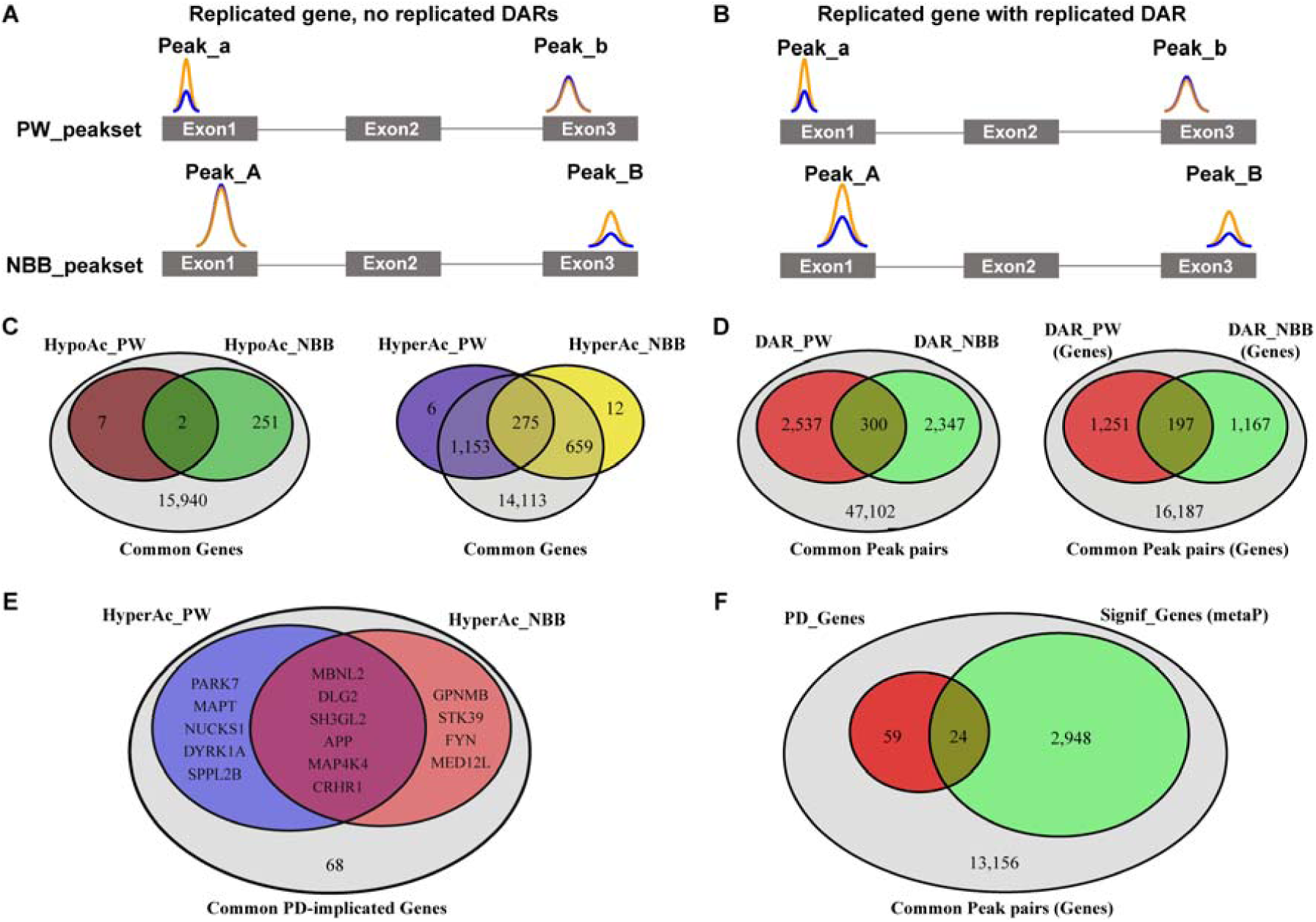
Replicated genes and replicated DARs. **A**,**B.** Schematic representation of two levels of replication between the PW and NBB cohorts. H3K27ac peaks are illustrated by colored lines, representing control (blue) and PD (orange) samples, with the height of the peak representing the peak intensity. Shown is a hypothetical gene X, with exons represented as grey boxes and introns represented by lines. In both cases, the gene has two pairs of common peaks: Peak_a/A, mapped to the first exon of the gene and Peak_b/B, mapped to the third exon of the gene. **A.** Altered H3K27 acetylation occurs at different regions of gene X in each cohort. Thus, while gene X harbors altered H3K27 acetylation in both cohorts (i.e. gene X is a replicated gene), it does not harbor replicated DARs. **B.** Altered H3K27 acetylation occurs at the same region of gene X in both cohorts. In this case, gene X is a replicated gene with a replicated DAR. **C.** Venn diagram showing the number of replicated genes (with or without replicated DARs) between the two cohorts, out of all genes represented in our ChIP-seq data from both cohorts. **D.** Venn diagrams showing the number of replicated DARs between the two cohorts, out of all common peak pairs (left) and their corresponding annotated genes (right). **E.** Venn diagram showing the overlap of PD-implicated genes with DARs between cohorts. All DARs exhibited hyperacetylation. **F.** Venn diagram showing the number of PD-implicated genes with adjusted metaP < 0.05.

### Meta-analysis of ChIP-seq data from PW and NBB cohorts

In order to perform meta-analysis of the ChIP-seq data we first harmonized the data from the two cohorts by pairing peaks with the same structural and functional genomic annotation (e.g. promoter, exon, UTR). These peaks are henceforth referred to as *“*common peaks*”*. For each common peak pair, we then calculated the Fisher’s meta p-value (metaP). A toy example of a common peak pair is shown in Fig. 4A,B.

In total, the harmonization process resulted in 52,286 pairs of common peaks. Out of these, 7,288 (mapped to 2,972 genes) had adjusted metaP < 0.05 (Supplementary Table S7, Fig. 4D,F). Moreover, 300 pairs represented peaks defined as DARs in both cohorts (Fig. 4D, hypergeometric p-value = 1.16 x 10^−3^). These 300 common peak pairs, mapped to 197 genes, are henceforth referred to as “replicated DARs”. Nearly all (295/300) replicated DARs were hyperacetylated in both cohorts and only a single DAR, mapped to the *PTPRH* gene, was hypoacetylated. Genes annotated to the top replicated DARs, ranked based on their adjusted metaP, are provided in Table 2.

**Table 2.** Genes annotated to the top replicated DARs.

| Symbol | Region Annotation | CHR | Region Location | Peak Location (PW, NBB) | MetaP | LFC PW, NBB | PeakName PW, NBB | Proportion Significant |
| --- | --- | --- | --- | --- | --- | --- | --- | --- |
| Common hypoacetylated regions |  |  |  |  |  |  |  |  |
| *PTPRH* | introns | 19 | 55718568-55720750 | 55720288-55722414, 55717912-55719103 | 1.7E-07 | -0.81, -0.94 | Peak_20188, Peak_15767 | 3<sup>p</sup>/4(0.8) |
| *JUP* | introns | 17 | 39681526-39923630 | 39805764-39808133, 39807092-39808006 | 1.1E-05 | -0.43, -0.99 | Peak_53983, Peak_81536 | 5<sup>p</sup>/10<sup>p</sup>(0.5) |
| Common hyperacetylated regions (top 15) |  |  |  |  |  |  |  |  |
| *DLG2\** | exons | 11 | 83874504-83874554 | 83873551-83879844, 83873666-83877544 | 7.7E-12 | 0.68, 1.05 | Peak_21808, Peak_40741 | 30<sup>p</sup>/49<sup>p</sup>(0.6) |
| *KIT* | exons | 4 | 55604595-55606881 | 55606747-55609274, 55606782-55609553 | 8.1E-11 | 0.63, 1.26 | Peak_21777, Peak_37103 | 2/3<sup>p</sup>(0.7) |
| *CNTNAP2* | exons | 7 | 147336198-147336398 | 147333973-147338260, 147333944-147337213 | 6.2E-09 | 0.51, 1.07 | Peak_44999, Peak_47588 | 11/18<sup>p</sup>(0.6) |
| *RNF216* | introns | 7 | 5681008-5692043 | 5679685-5685127, 5688370-5705729 | 1.2E-08 | 0.6, 0.53 | Peak_38631, Peak_12618 | 11<sup>p</sup>/16(0.7) |
| *FRMD5* | introns | 15 | 44216508-44393656 | 44260324-44263821, 44244249-44247825 | 1.6E-08 | 0.68, 0.67 | Peak_18018, Peak_18744 | 12<sup>p</sup>/17<sup>p</sup>(0.7) |
| *SOX2-OT* | introns | 3 | 181328718-181417385 | 181358880-181360150, 181341154-181348014 | 1.7E-08 | 0.55, 0.91 | Peak_43667, Peak_31549 | 9/12(0.8) |
| *MAGI2* | 1to5kb | 7 | 78120338-78124337 | 78121257-78128843, 78115190-78121170 | 2.2E-08 | 0.4, 0.93 | Peak_18816, Peak_24042 | 9<sup>p</sup>/23(0.4) |
| *PPP2R2B* | introns | 5 | 146080706-146460621 | 146383635-146384203, 146459075-146469815 | 2.8E-08 | 1.3, 0.52 | Peak_135972, Peak_17645 | 17<sup>p</sup>/27<sup>p</sup>(0.6) |
| *KCNH1* | introns | 1 | 210948887-210970849 | 210966438-210969215, 210946491-210950181 | 3.8E-08 | 0.55, 1.0 | Peak_51838, Peak_38212 | 9/18<sup>p</sup>(0.5) |
| *CDH1* | introns | 16 | 68835797-68842326 | 68840159-68841130, 68782065-68837136 | 4.8E-08 | 0.84, 0.37 | Peak_66179, Peak_3389 | 5/7<sup>p</sup>(0.7) |
| *MAP4K4\** | introns | 2 | 102315001-102407181 | 102330963-102334475, 102367013-102375949 | 5.2E-08 | 0.54, 0.62 | Peak_30093, Peak_9009 | 10<sup>p</sup>/24<sup>p</sup>(0.4) |
| *METTL24* | introns | 6 | 110644076-110729536 | 110656829-110661962, 110685537-110688887 | 5.4E-08 | 0.48, 0.97 | Peak_22739, Peak_27591 | 8/13<sup>p</sup>(0.6) |
| *PIP4K2A* | introns | 10 | 22898647-23003111 | 22934570-22952399, 22957818-22962517 | 5.6E-08 | 0.58, 0.52 | Peak_22480, Peak_27954 | 9<sup>p</sup>/18<sup>p</sup>(0.5) |
| *SIK3* | introns | 11 | 116827781-116968858 | 116825819-116835556, 116846923-116884577 | 5.8E-08 | 0.6, 0.49 | Peak_21504, Peak_11226 | 13<sup>p</sup>/16<sup>p</sup>(0.8) |
| *DOCK10* | introns | 2 | 225796386-225906968 | 225798681-225800264, 225845772-225849871 | 8.1E-08 | 0.61, 0.77 | Peak_94440, Peak_22866 | 6/10<sup>p</sup>(0.6) |
| Common hyperacetylated regions (additional GWAS hits) |  |  |  |  |  |  |  |  |
| *CRHR1\** | introns | 17 | 43707525-43750192 | 43726544-43728643, 43741886-43747152 | 2.0E-07 | 0.43, 0.83 | Peak_74390, Peak_30944 | 6/14<sup>p</sup>(0.4) |
| *MBNL2\** | introns | 13 | 97873814-97927885 | 97919075-97922481, 97906667-97916758 | 1.2E-05 | 0.44, 0.44 | Peak_29656, Peak_16279 | 14<sup>p</sup>/22<sup>p</sup>(0.6) |

\* - Previous association with PD based on GWAS; Region Location – genomic location (on the indicated chromosome) of the annotated region; Peak location – genomic location of the relevant peak from each cohort (upper – PW, lower - NBB); LFC – log fold change compared to controls; Peak Name – the peak identifier in each cohort; Proportion significance – DARs/total common regions annotated to the gene; <sup>p</sup>- peaks annotated to promoter region of the gene.

### Altered H3K27 acetylation has a predilection for genes with an established link to PD

We next wondered whether genes with an established involvement in PD are over-represented among DAR-harboring genes. To assess this, we compiled a list of genes that fulfilled at least one of the following criteria: (a) are associated with PD based on the most recent and largest GWAS metaanalysis^4^, (b) are implicated in monogenic PD^37,38^, or (c) encode proteins with a central role in PD-related neuropathology^2,39,40^. Among the 92 genes meeting these criteria (Supplementary Table S8), 83 could be assessed in ChIP-seq data from both cohorts and are henceforth referred to as “PD-implicated genes”. Out of these, six genes: *DLG2, MAP4K4, CRHR1, MBNL2, SH3GL2* and *APP*, harbored DARs in both cohorts (hypergeometric p-value = 2.4 x 10^−3^, Fig. 4E). In addition, 24/83 PD-implicated genes had adjusted metaP < 0.05 (hypergeometric p-value = 6.1 x 10^−3^, Supplementary Table S8, Fig. 4F). These included genes with key-roles in the pathogenesis of idiopathic and/or monogenic PD: *SNCA, MAPT, APP, PRKN* and *PARK7*. Interestingly, the significantly hyperacetylated peak in *SNCA* was in an enhancer region of the gene previously shown to be affected by both genetic variation^41^ and drug exposure associated with PD^42^ (Fig. 5A).

**Fig. 5:**
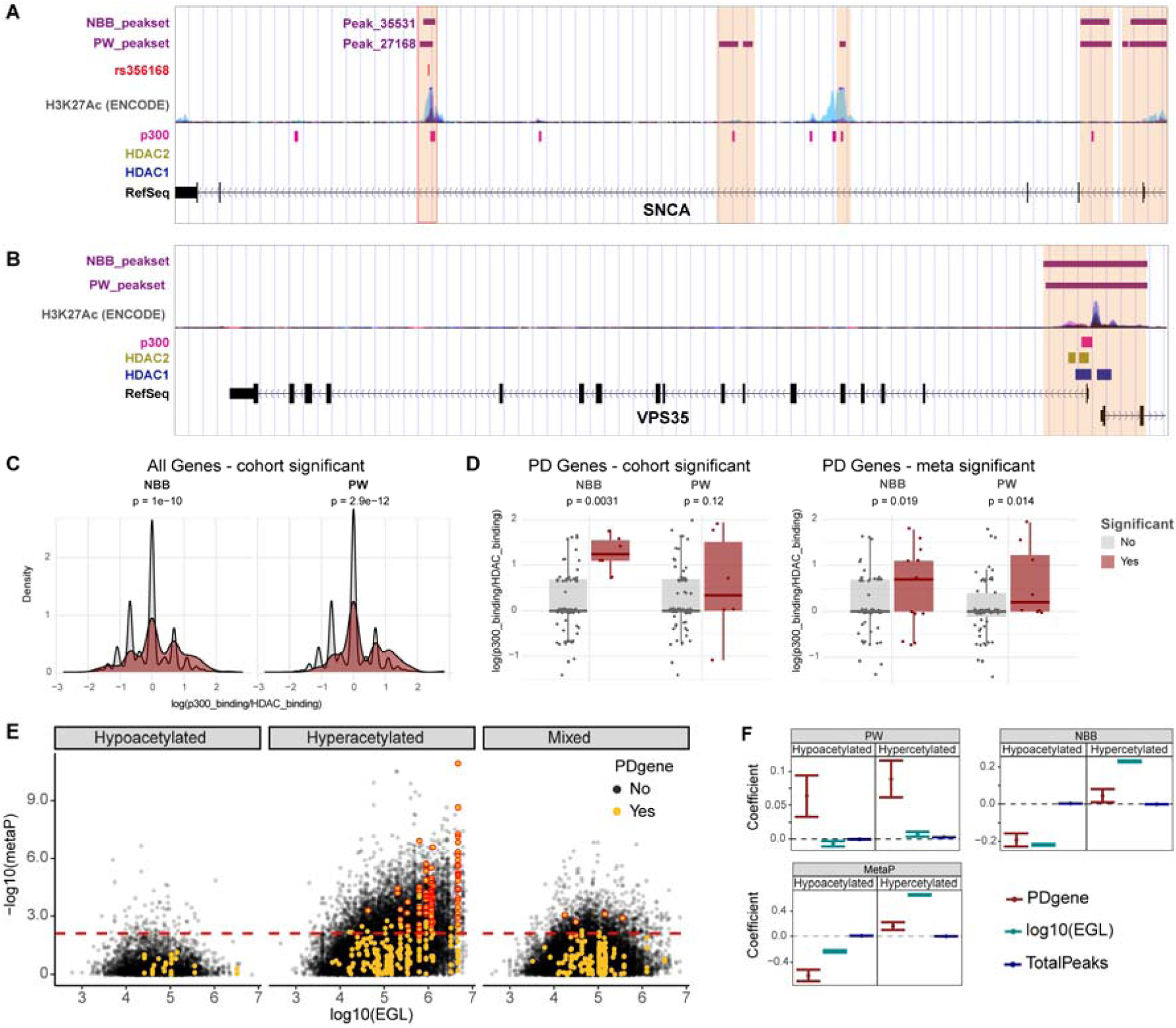
H3K27 acetylation is associated with the relative proportion of p300 and non-SIRT1 HDAC binding and overrepresented in PD-implicated genes. **A**,**B.** Examples of two PD_implicated genes, *SNCA* (**A**) and *VPS35* (**B**). *SNCA* harbors a DAR (Peak_35531/27168 in PW/NBB respectively) in an enhancer element whose H3K27 acetylation status is associated with both genetic variation (SNP rs356168) and drugs influencing the risk of PD. *VPS35* does not exhibit DARs in our data. Purple bars indicate H3K27ac peaks in the PW and NBB peaksets. H3K27ac ChIP-seq data from ENCODE is shown for reference. Pink, blue and yellow bars indicate the binding sites of p300, HDAC1 and HDAC2 respectively, based on ENCODE ChiP-seq data wgEncodeRegTfbsClusteredV3.bed. **C**,**D** Difference in the relative number of p300 and non-SIRT1 HDAC bindings sites between significantly hyperacetylated and non-significantly hyperacetylated peaks for all (**C**, Kolmogorov–Smirnov test) or only PD_implicated (**D**, two-sided Wilcoxon rank sum test) genes. **E.** Hyperacetylated – regions showing hyperacetylation in both cohorts; Mixed – regions with opposite acetylation trends in the two cohorts. The dashed line represents the -log10 of the highest metaP corresponding to adjusted metaP < 0.05. Multiple common regions within the same gene are shown. Highlighted are regions identified as DAR in at least one of the cohorts within PD-implicated genes. metaP – Fisher’s meta-p value of the corresponding two peaks from PW and NBB cohorts. **F.** Association between -log(p) and each of the indicated covariates based on linear model. Shown are the coefficients and 95% confidence intervals. The results are shown for the common regions (metaP) and each of the cohorts separately. Peaks annotated to miRNAs and snRNAs were excluded from the analysis. PDgene – a gene is a PD-implicated gene. EGL – effective gene length. TotalPeaks – total number of common regions or peaks annotated to the gene.

### H3K27 hyperacetylated regions are associated with an increased ratio of p300 binding sites relative to non-SIRT1 HDAC binding sites

Acetylation of H3K27 is largely mediated by the histone acetyl transferase p300^43^, whose activity is suppressed by SIRT1^44^. Thus, we reasoned that if H3K27 hyperacetylation in PD is indeed mediated by decreased SIRT1 activity, it should be associated with regions exhibiting increased binding of p300 and decreased binding of non-SIRT1 HDACs. To test this, we integrated our data with the relevant ChIP-seq data derived from the ENCODE project^45^ (see Methods). Specifically, we quantified the number of p300 bound regions (p300_binding) or non-SIRT1 HDAC bound regions (HDAC_binding) from ENCODE that overlap with H3K27ac peaks in our data and assessed the association of the two measures with H3K27 hyperacetylation level of each region. In both cohorts, the ratio of the of p300 binding sites over non-SIRT1 HDAC binding sites was significantly higher in regions overlapping with hyperacetylated DARs compared to non-DARs. Notably, this was also true when restricting the analyses to the PD-implicated genes (Fig. 5C,D). Linear model analysis showed positive association of H3K27 hyperacetylation with p300_binding (PW: β = 0.02, p < 2 x 10^−15^; NBB: β = 0.09, p < 10^−26^) and negative association with HDAC_binding (PW: β = -0.07, p < 10^−26^; NBB: β = -0.14, p < 10^−26^). Similar results were obtained when peaks were collapsed to a gene level (Supplementary Table S9).

### Differential H3K27 acetylation is associated with effective gene length

We noted that many of the genes with DARs, including PD-implicated genes, are among the longest in the genome. We reasoned that this may be because longer genes have higher histone binding capacity and therefore are more likely to show a differential signal in the event of a generalized H3K27 hyperacetylation. To test this hypothesis, we used linear models to assess whether H3K27ac, defined as -log10(metaP), is associated with PD-implicated genes while accounting for the effective gene length (EGL), defined as the sum of all regions annotated to a gene (see Methods). The total number of common regions in a gene was included as a covariate in the model. For hyperacetylated regions, this analysis indicated a significant positive association between altered acetylation and PD-implicated genes (p = 1.0 x 10^−7^) as well as EGL (p < 2.0 x 10^−26^). For the hypoacetylated regions, however, the analysis indicated a significant negative association with both PD-implicated genes (p < 2.0 x 10^−26^) and EGL (p < 2.0 x 10^−26^). Analysis of each cohort independently corroborated these findings, although the negative association with PD-implicated genes for the hypoacetylated regions was only significant in the NBB cohort (Fig. 5B, Supplementary Table S10).

### Promoter H3K27 acetylation is decoupled from gene expression in PD

While it is well accepted that modifications of H3K27ac are predictive of gene expression levels^46^, it was previously shown that widespread histone hyperacetylation induced by potent HDAC inhibition does not lead to major changes in gene expression^47,48^. To investigate whether a similar decoupling occurs in the PD brain, we analyzed RNA sequencing (RNA-seq) data from the PFC of the same individuals included in the ChIP-seq analysis (Supplementary Table 1, Supplementary Methods). For each gene, we then calculated the Pearson’s correlation between the level of its expression and promoter H3K27ac across controls or individuals with PD.

In the control group from both cohorts, most genes exhibited the expected positive correlation between promoter H3K27ac and expression. In the PD group, however, the distribution of correlations was centered closer to 0, regardless of the absolute correlation threshold chosen (Fig. 6A-E). The deviation between the two groups progressively increased as only genes with higher correlation between promoter H3K27ac and expression were included in the analysis (Fig. 6B-E). Importantly, similar results were obtained based on RLE-normalized ChIP-seq data (Supplementary Fig. S8), indicating that the observed decreased correlation between H3K27ac and transcription levels in PD was not driven by our normalization method. These results suggest that the coupling between promoter H3K27 acetylation state and proximal gene transcription is impaired in the PFC of individuals with PD.

**Fig. 6:**
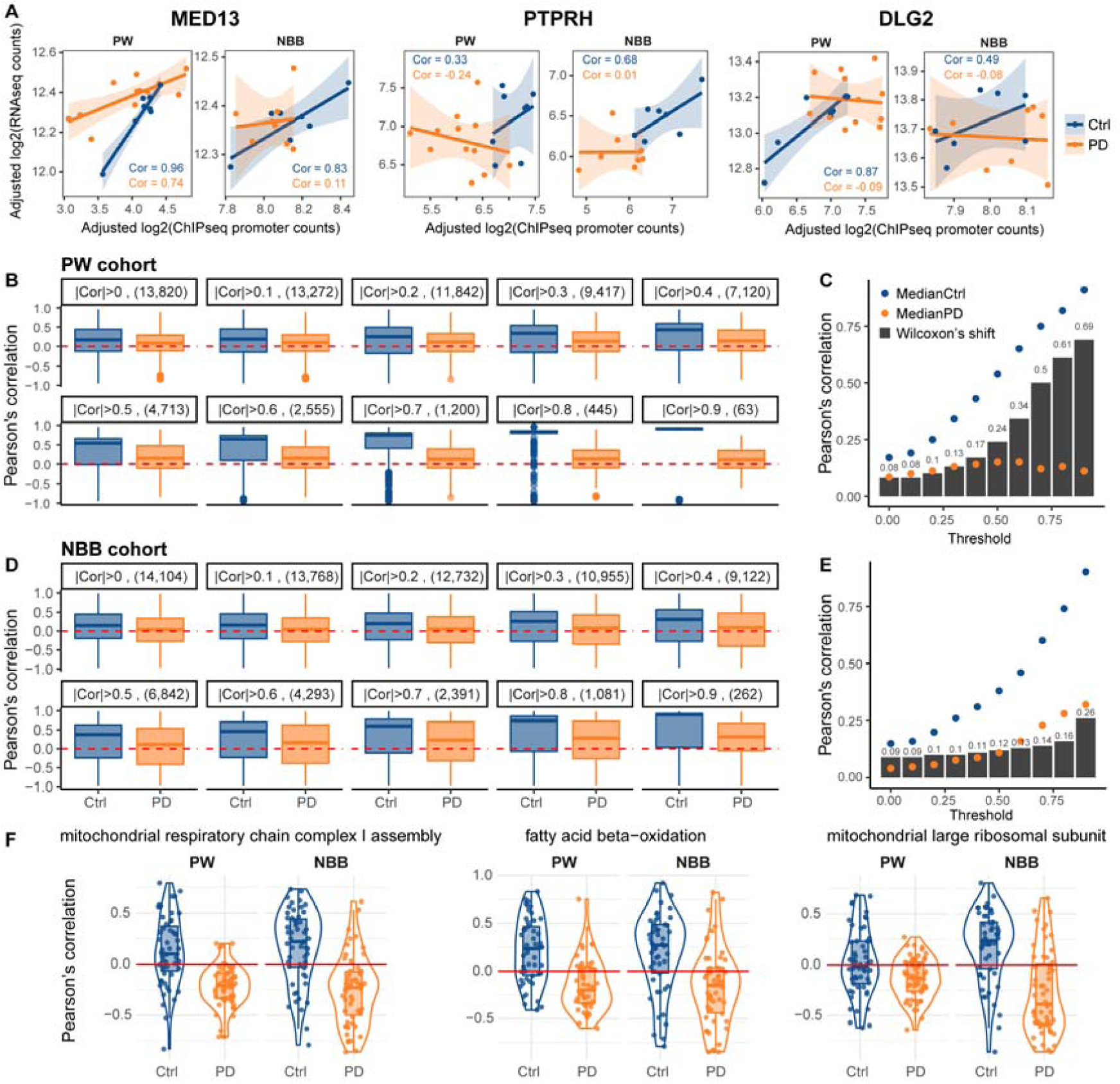
Decreased correlation between promoter H3K27 acetylation state and gene expression in PD. Pearson’s correlation between the adjusted promoter H3K27ac counts and the adjusted RNA-seq counts was calculated for each gene across control or PD subjects in each cohort separately. **A.** Representative scatter plots showing the adjusted promoter H3K27ac counts (X-axis), the adjusted RNA-seq counts (Y-axis) and the calculated Pearson’s correlation for three genes: *MED13, PTPRH* and *DLG2* in control (blue) or PD (orange) individuals from the PW or NBB cohorts. Each dot represents an individual. Cor - Pearson’s correlation. **B-E**, The distribution of correlations for all compatible genes was compared between the groups for various thresholds of minimal absolute correlation in either of the groups. Results are shown for PW (**B**,**C**) and NBB (**D**,**E**) cohorts. The number of genes that remained after applying the correlation threshold is shown in parentheses. The correlations remained closely distributed around 0 in PD subjects from both cohorts, regardless of the correlation threshold. **C**,**E**, Median correlations in each group and the calculated Wilcoxon’s delta shift for the different correlation thresholds. **F.** Nucleus-encoded mitochondrial genes exhibit negative correlation between promoter H3K27 acetylation and gene expression. Shown are examples for three mitochondria related GO terms.

Functional enrichment analysis based on the decoupling level of the genes, indicated that nucleus-encoded mitochondrial genes are the most decoupled genes in both cohorts (Supplementary Table S11). Moreover, closer examination of these genes revealed that most of them exhibit negative correlation between the expression level and promoter H3K27 acetylation (Fig. 6F, Supplementary Table S11).

## Discussion

### Aberrant histone acetylation occurs genome-wide in the PD brain

We report increased acetylation of multiple sites on histones H2B, H3 and H4 with the strongest change observed for H3K27, a marker of active promoters and enhancers with a fundamental role in regulating gene expression^49,50^. Genome-wide H3K27 hyperacetylation in PD is supported by multiple levels of evidence in our study: (a) immunoblot analyses showed a highly significant increase in the acetylated fraction of H3K27 across three brain regions in PD. Importantly, we show that this hyperacetylation is not driven by underlying changes in cell composition and is unlikely to be the result of anti-Parkinson medication. (b) ChIP-seq analysis, replicated in two independent cohorts, showed that the total number of peaks, percentage of unique peaks, genome coverage and the total number of RiPs are all increased in the PD samples. The analysis further revealed that increased H3K27ac occupancy has a genome-wide distribution. (c) We found a strong association between differential H3K27 acetylation and the effective length of the targeted gene. This association was positive for hyperacetylated regions and negative for hypoacetylated regions, in line with a hyperacetylation-predisposing cellular environment in PD.

### H3K27 hyperacetylation is likely to be induced by altered sirtuin activity

We hypothesized that H3K27 hyperacetylation in PD may be mediated by altered sirtuin activity, due to decreased NAD^+^/NADH ratio^15,18,51^ resulting from complex-I deficiency, a pathological hallmark of the PD brain^13,14^. Hyperacetylation of several histone sites, including H3K27, was indeed observed in animal and cell studies, following chemical complex-I inhibition with PD-associated neurotoxins^10,52^. In support of our hypothesis, we found increased levels of SIRT1 and SIRT3 proteins in PD samples and no change in SIRT2 quantity, or in the acetylation state of its substrate, α-tubulin. This is consistent with the notion that SIRT1 and SIRT3, but not SIRT2 activity are susceptible to changes in physiological NAD^+^ concentrations^51^. In line with our findings, reduced sirtuin activity accompanied by increase in their protein levels was reported in iPSC-derived dopaminergic cells carrying a *LRRK2* mutation^53^. Moreover, complex-I deficiency, decreased NAD^+^/NADH ratio, decreased sirtuin activity (accompanied by increased protein levels), and increased histone acetylation have all been shown to occur with aging^5,17,20,22,23^, which is the strongest known risk factor for PD.

The role of SIRT1 in mediating H3K27 hyperacetylation in PD is further supported by the observation that hyperacetylation is more pronounced among regions with high p300/non-SIRT1 HDAC binding site ratio. This is because decreased SIRT1 activity increases p300-mediated H3K27 acetylation, and can be compensated by other HDACs. Thus, H3K27 hyperacetylation induced by decreased SIRT1 activity will be more pronounced in regions exhibiting multiple p300 bindings sites and fewer non-SIRT1 HDAC binding sites.

### H3K27 hyperacetylation is over-represented in PD-implicated genes

While H3K27 hyperacetylation appears to be a genome-wide phenomenon in the PD brain, it also shows a highly significant predilection for genes implicated in PD. Meta-analysis of our two cohorts identified H3K27 hyperacetylated regions in 24/83 genes linked to idiopathic and/or monogenic forms of PD (Supplementary Table S8). These include 21 genes associated with PD by GWAS^4^, genes encoding proteins involved in PD pathology (*SNCA, MAPT, APP*)^2,39,40^ and genes in which mutations cause Mendelian PD (*SNCA, PRKN, PARK7*)^54,55^. In addition, H3K27 hyperacetylation was observed in genes associated with more complex forms of parkinsonism and/or vulnerability to PD-related pathology such as *FBOX7*^56^ and *POLG*^37^.

Several mechanisms may underlie the observed over-representation of PD-implicated genes. First, it is plausible that H3K27 acetylation at promoter and/or enhancer regions of these genes is modulated by genetic variation and/or environmental factors associated with PD. For instance, the H3K27 acetylation state of the *SNCA* enhancer region, significantly hyperacetylated in our data, was shown to be affected by both genetic variation^41^ and drug exposure associated with PD^42^. Second, it is likely that pathogenic processes involved in PD induce the expression of genes implicated in the disease. For example, the expression levels of *PRKN* and *PARK7* increase in response to oxidative stress^57–60^. Since, the regulation of transcription often involves histone acetylation, impairment of the deacetylation machinery, would over time lead to excessive H3K27 acetylation of these genes. Finally, by relaxing chromatin structure, H3K27 hyperacetylation in genomic regions associated with PD, predisposes these areas to somatic mutations, including genomic rearrangements^61^. As previously suggested^62^, this could act synergistically with other predisposing factors, such as age and oxidative injury, eventually leading to PD-specific pathology.

### H3K27 hypoacetylation in the *PTPRH* gene may contribute to PD pathophysiology

Considering the general H3K27 hyperacetylation in the PD brain, the robust hypoacetylation of H3K27 in the *PTPRH* gene, encoding protein tyrosine phosphatase receptor type H, is of particular interest. While little is known about the physiological function of *PTPRH*, it was previously found to be enriched in loss-of-function mutations in PD^35^. Furthermore, it was shown that RNAi-mediated downregulation of the *PTPRH* homolog in *Drosophila* enhances α-synuclein neurotoxicity^35^. Our finding of selective H3K27 hypoacetylation of the *PTPRH* region in PD, corroborates the hypothesis that decreased function of this gene may contribute to the pathophysiology of PD.

### Decoupling between promoter H3K27 acetylation and gene expression in the PD brain

The attenuated correlation between promoter H3K27 acetylation and gene expression, is intriguing, since it indicates that a fundamental mechanism of gene regulation is defective in the PD brain. Furthermore, since histone acetylation is tightly linked to the metabolic state of the cell^51^, this decoupling suggests that cellular responses to metabolic stress may be severely compromised in PD. This putative compromised response to metabolic stress is further supported by the finding that, in both cohorts, the highest degree of decoupling is observed among nuclear-encoded mitochondrial genes. Moreover, the negative correlation between H3K27 acetylation and gene expression of these genes supports the involvement of sirtuins, in particular SIRT1, in the pathophysiology of PD. This is because in addition to histones, SIRT1 deacetylates and activates PGC-1α, the key regulator of mitochondrial biogenesis^18,51,63^. Thus, decreased SIRT1 activity would simultaneously lead to increased histone acetylation and reduced expression of nuclear-encoded mitochondrial genes, resulting in negative correlation between their expression level and promoter H3K27 acetylation state. Moreover, decreased expression of mitochondrial genes would initiate a vicious circle by decreasing NAD^+^/NADH ratio and further reducing SIRT1 activity. In conclusion, we show that H3K27 acetylation, a fundamental mechanism modulating gene expression, is severely dysregulated in the PFC of individuals with idiopathic PD. The highly significant concordance between the two cohorts and the clear predilection for genes strongly linked to PD, validate our findings and suggest that epigenetic dysregulation plays an important role in the pathogenesis of the disease. Importantly, since histone acetylation and NAD metabolism can be pharmaceutically modulated, our results open the possibility that agents targeting these biological processes may have therapeutic potential for PD and should be further explored.

## Materials and Methods

### Cohorts and tissue

Brains were collected at autopsy and split sagittally along the corpus callosum. One hemisphere was fixed whole in formaldehyde for ∼2 weeks, and the other coronally sectioned and snap-frozen in liquid nitrogen. Prefrontal cortex (Brodmann area 9) was obtained from two independent cohorts. The discovery cohort (ParkWest, or PW cohort) comprised individuals with idiopathic PD (n = 17) from the ParkWest study, a prospective population-based cohort which has been described in detail^12^, and neurologically healthy controls (n = 15) from the Neuromics Biobank for Aging and Neurodegeneration. For a subset of the PW cohort (PD: n =7, Control: n = 7), tissue samples from the striatum (putamen) and cerebellar cortex were also included. All individuals of the PW cohort had undergone whole-exome sequencing^64^ and individuals with pathogenic mutations in genes implicated in Mendelian PD and other monogenic neurological disorders had been excluded. The replication cohort, comprised individuals with idiopathic PD (n = 10) and neurologically healthy controls (n = 11) of Dutch origin, from the Netherlands Brain Bank (https://www.brainbank.nl/). Demographic information and experimental allocation of all samples is shown in Supplementary Table S1. For each experiment, there were no significant differences for sex, age, or postmortem interval (PMI) between the compared groups. A complete list of the medication used by the individuals with PD during at least the last 12 months before death can be found in Supplementary Table S1. Individuals with PD fulfilled the National Institute of Neurological Disorders and Stroke^65^ and the U.K. PD Society Brain Bank^66^ diagnostic criteria for PD at their final visit. All cases showed neuropathological changes consistent with PD.

### Immunoblotting

Three independently dissected samples of brain tissue (replicates) were analyzed from each individual. Immunoblots were repeated at least 2 times for each marker. To normalize the histone acetylation signal to histone quantity, each acetylation blot was stripped twice. Complete stripping was confirmed by incubation of the membranes with secondary antibodies only and ECL. To exclude potential bias due to incomplete stripping, we performed additional experiments where acetylated and total histone markers for each histone modification were assessed in parallel on different blots. To estimate the acetylated fraction of each histone marker, we calculated the signal ratio of acetylated marker to total histone protein. This ratio is referred to as the “acetylation fraction” for each histone marker. Detailed description is provided in the Supplementary material, methods section. Comparison of immunoblot densitometric values between groups was performed using linear regression for panacetylation and linear mixed-models for all other markers. Linear mixed-models analysis was chosen in order to account for intraindividual random variability. Subjects were specified as random effects and disease state, sex, age of death, PMI and oligodendrocyte MSP were used as fixed effects. The error distribution was assumed normal, and solution to the mixed models was obtained using the maximum likelihood estimation method. The mixed-models analyses were implemented in SPSS version 22.0.0.

### Chromatin Immunoprecipitation sequencing (ChIP-seq)

Approximately 200mg of frozen PFC tissue per sample was used for ChIP-seq. Briefly, the tissue was crosslinked using formaldehyde, lysed using a Dounce homogenizer and sonicated to obtained chromatin fragment of 300-500 bp. Chromatin was immunoprecipitated using an antibody against acetylated H3K27. Protein-DNA complexes were recovered using protein A agarose beads, de-crosslinked and DNA was purified by phenol-chloroform extraction and ethanol precipitation. Following quantitative PCR, libraries were built and sequenced on Illumina’s NextSeq 500 (75 base pair reads, single end). Detailed description is provided in the Supplementary material, methods section.

### ChIP-seq quality control and filtering

Raw FASTQ files were trimmed using Trimmomatic version 0.38^67^ with options ILLUMINACLIP:TruSeq2-SE.fa:2:30:10 LEADING:3 TRAILING:3 SLIDINGWINDOW:4:15 MINLEN:25. Assessment of FASTQ files was carried out with FastQC version 0.11.8 [http://www.bioinformatics.babraham.ac.uk/projects/fastqc] before and after trimming. Reads were aligned to the GRCh37 human genome using bwa version 0.7.17^68^. Reads with mapping quality < 30 were discarded using samtools version 1.9 [http://www.htslib.org/] and duplicated reads removed using MarkDuplicates from Picard Tools version 2.18.27 [http://broadinstitute.github.io/picard/]. BAM files were converted into BED files and sorted.

### Peak calling and read counting

Peak calling procedure was carried out in two complementary approaches: 1) **Per group (PD or controls):** the peaks are called separately for each group, aggregating the relevant samples and inputs. While this approach is not optimal for differential peak calling analysis as it requires harmonization of peaks between the two groups, it can be used to compare the total genome coverage as well as unique H3K27ac genomic regions in each group. 2) **All samples combined:** the peaks are called by aggregating all samples and input controls. This approach does not allow identification of unique genomic regions however, it provides a better identification of consensus peaks, and thus, was used for differential peak analysis. The resulting peak-sets were filtered using the ENCODE black listed regions and only peaks within canonical chromosomes were kept for downstream analyses. Sample-specific enrichment on the identified peaks was performed using featureCounts version 1.6.4^69^ with default parameters. For both the individual group and all samples combined approaches, peaks with non-adjusted p-value > 10^−7^ were excluded from the analysis. Reads inside peaks (RiP) were quantified using “featureCounts” program from the “subread” package v2.0.0. Additional details are provided in the Supplementary material, methods section

### Calculation of Marker Site Profiles (MSP)

In order to identify brain cell-type specific H3K27ac regions we first analyzed H3K27ac ChIP-seq data from NeuN^+^ and NeuN^-^ brain cells^25^. Cell-type based broadPeaks (CellType_peak-set), count matrixes and metadata files were downloaded from https://www.synapse.org/#!Synapse:syn4566010. Differential regions between NeuN^+^ and NeuN^-^ cells were calculated using “DESeq2” R package^70^, including chromatin amount, library batch, sex, hemisphere, age and pH as covariates in the model. Peaks were defined as cell-type specific differentially acetylated regions (DARs) if they met the following criteria: a) |fold change| > 4 and b) mean count > 1000. Peaks were annotated to all genes for which they intersected a region between 5kb upstream from their transcription start site (TSS) to the end of their 5’UTR using “annotatr” R package^71^, based on UCSC hg19 genome assembly. Genes with DARs were then itersected with expression-based cortical marker gene lists based on NeuroExpresso database^27^. The DARs were next reassigned to specific cell-types if they were annotated to genes defined as cell-type specific based on NeuroExpresso. For example, all DARs annotated to *MBP*, defined as oligodendrocyte marker gene based on NeuroExpresso, were defined as oligodendrocyte marker sites (MSS). In the next step, we quantified the reads from our samples in the CellType_peak-set. The corresponding RiP were then converted to counts per million (CPM) and transformed to log2(CPM + 1). Next, for each cell-type specific MSS, we performed principal component analysis based on the relevant peaks as described in^27^. Marker Site Profiles (MSP) were defined as the scores of the samples in the first principal component, transformed to [0,1] range for visualization purposes. Detailed description of the analysis is provided in the Supplementary material, methods section.

### Identification of differentially acetylated regions

Differential peak analysis was performed using “nbinomWaldTest” function from “DESeq2” R package. Dispersions were calculated by using “estimateDispertions” function from “DESeq2” R package, by setting the fitType parameter to “local”. Normalization factors calculated by the “estimateSizeFactors” function were replaced with the housekeeping normalization factor based on our manually selected housekeeping gene set (*ACTB, GAPDH, UBC*). Peak significance was determined using DESeq2 “results” function, setting the cutoff for adjusted p-value to 0.05, and independentFiltering=T. Detailed description of the analysis is provided in the Supplementary material, methods section.

### Association of H3K27 hyperacetylation with p300 and non-SIRT1 HDAC binding

ChIP-seq data (wgEncodeRegTfbsClusteredV3.bed) was obtained from the Encyclopedia of DNA Elements (ENCODE) project^45^. The data was available for p300 and the following HDACs: HDAC1, HDAC2, HDAC6, HDAC8 and SIRT6 We first quantified the number of binding sites for each protein in our peaks using the “findOverlaps” function from the “GenomicRanges” R package^72^. For each peak, the HDAC binding sites were then summed up to represent the non-SIRT1 HDAC_binding. Peaks that had no binding sites for either of the proteins were excluded from downstream analyses. For the gene level analysis, the number of binding sites in each peak was collapsed to a gene level.

Difference between DARs and non DARs was assessed by comparing the value of 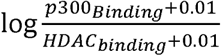 of the peaks using Kolmogorov-Smirnov test (all genes) or Wilcoxson’s rank sum test (PD_implicated genes) using “ks.test” and “wilcox.test” functions from R “stats” package. For linear model, we first obtained one-sided hyperacetylation p-value (p_hyper_) for each peak as followed: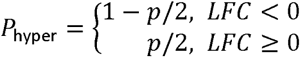. The association of hyperacetylation with p300_binding and HDAC binding was then assessed by the linear model: −*log*10(*p*_*hyper*_)∼ *p*300_*binding*_ + *HDAC*_*binding*_ + *PDgene* where PDgene indicates whether a peak is annotated to PD_implicated gene or not.

### Association of PD_implicated genes and EGL with altered H3K27 acetylation

Effective gene length (EGL) was defined as sum of length of all regions annotated to the gene (including 1Kb upstream to TSS). We assessed the association between altered H3K27 acetylation defined as -log10(metaP) with being annotated to a PD_implicated gene (PDgene), EGL and the total number of common regions aligned to the gene using a linear model. The total number of common regions in a gene was included as a covariate in the model. Namely, for a common region i, annotated to gene j, the -log(metaP) value was modeled as: −*log*10(*metaP*_*ij*_) ∼ *PDgene*_*j*_ + *log*10(*EGL*_*j*_) + ∑*CommonRegions*_*j*_. For the analysis was performed in each cohort separately, metaP was replaced with cohort-specific p-value, and the total number of common regions was replaced with the total number of regions (peaks) annotated to the gene in the cohort-specific peak-set.

### RNA-seq analysis

Total RNA was extracted from prefrontal cortex tissue homogenate for all samples for which ChIP-seq was performed and sequenced following ribosomal RNA depletion A detailed description of the RNA-seq analysis is provided in the Supplementary material, methods section and is reported elsewhere^73^. Relative cell abundance was estimated using cell-type specific marker genes, as previously described^27,74^. Since significant group effect was found for oligodendrocytes and microglia estimates in PW (Wilcoxon’s p = 2.6 × 10^−4^) and NBB (Wilcoxon’s p = 0.009) cohorts respectively, these cell types were included at covariates in the analysis of both cohorts. Count matrix and the code necessary to reproduce these results are available on https://git.app.uib.no/neuromics/cell-composition-rna-pd.

### Promoter H3K27ac - expression correlation

ChIP-seq and RNA-seq counts were first adjusted to the relevant covariates (age, sex, PMI, batch, oligodendrocyte estimates and microglia estimates). The adjustment was done by re-fitting the counts using the coefficients calculated by DESeq2 R function with adjusted model matrix in which categorical covariates were set to 0, and the continuous covariates to the mean of the original values. Next, for each gene, Pearson’s correlation between the adjusted ChIP-seq/RNA-seq seq counts was computed using “cor.test” R function, in each group (controls or PD subjects) separately. Only peaks annotated to promoter regions of the genes were included in the analysis. When multiple promoter peaks were defined for a gene, the correlation was calculated for each promoter separately, and the highest correlation was kept. This step was performed independently for each group. Since not all genes are expected to exhibit a good correspondence between promoter H3K27ac state and expression level due to existence of multiple levels of regulation, we compared the distribution of correlations in each group for different thresholds of absolute correlation in either of the groups (e.g., cor > |0.1| means that only genes with absolute correlation > 0.1 in at least one of the groups are included in the analysis). Group comparison was performed using Wilcoxon rank sum test, using “wilcox.test” function from R “stats” function. Functional enrichment analysis was performed using the gene score resampling method implemented in the ermineR package version 1.0.1^75^, an R wrapper package for ermineJ^76^, setting the value of “BigIsBetter” to FALSE, with the complete Gene Ontology (GO) database annotation^77^. For the enrichment of genes based on the decoupling levels, gene scores were defined as Δ*Cor* = *Cor*_*PD*_ - *Cor*_*Control*_. For enrichment of genes based on the lowest correlation in PD group, gene scored were defined as the correlation values in the PD group.

### Data and code availability

The R code necessary to reproduce the results presented in the paper is available on https://github.com/ltoker/ChIPseqPD. Raw counts for the ChIP-seq analyses are available in the same repository (https://github.com/ltoker/ChIPseqPD/tree/master/data). Additional R code to reproduce the RNA-seq analyses is available on https://git.app.uib.no/neuromics/cell-composition-rna-pd. The corresponding raw data counts are included in the repository (https://git.app.uib.no/neuromics/cell-composition-rna-pd/tree/master/Data).

## List of abbreviations

ChIP-seq: chromatin immunoprecipitation sequencing
Ctrl: control
DAR: differentially acetylated region
EGL: effective gene length
GWAS: genome-wide association study
HAT: histone acetyltransferase
HDAC: histone deacetylase
L-DOPA: levodopa
mtDNA: mitochondrial DNA
MSPs: Marker Site Profiles
NAD: nicotinamide adenine dinucleotide
NBB: Netherlands Brain Bank
PD: Parkinson’s disease
PFC: Prefrontal cortex
PMI: Postmortem interval
PW: Park West
RiP: Reads in peaks
RLE: Relative Log Expression
SNc: substantia nigra pars compacta
TSS: transcription start site
WB: Western blot
WES: whole exome sequencing

## Acknowledgments

This work is supported by grants from The Research Council of Norway (ES633272) and Bergen Research Foundation (BFS2017REK05). The cell-type specific H3K27ac ChIP-seq data were generated as part of the PsychENCODE Consortium, supported by: U01MH103392, U01MH103365, U01MH103346, U01MH103340, U01MH103339, R21MH109956, R21MH105881, R21MH105853, R21MH103877, R21MH102791, R01MH111721, R01MH110928, R01MH110927, R01MH110926, R01MH110921, R01MH110920, R01MH110905, R01MH109715, R01MH109677, R01MH105898, R01MH105898, R01MH094714, P50MH106934, U01MH116488, U01MH116487, U01MH116492, U01MH116489, U01MH116438, U01MH116441, U01MH116442, R01MH114911, R01MH114899, R01MH114901, R01MH117293, R01MH117291, R01MH117292 awarded to: Schahram Akbarian (Icahn School of Medicine at Mount Sinai), Gregory Crawford (Duke University), Stella Dracheva (Icahn School of Medicine at Mount Sinai), Peggy Farnham (University of Southern California), Mark Gerstein (Yale University), Daniel Geschwind (University of California, Los Angeles), Fernando Goes (Johns Hopkins University), Thomas M. Hyde (Lieber Institute for Brain Development), Andrew Jaffe (Lieber Institute for Brain Development), James A. Knowles (University of Southern California), Chunyu Liu (SUNY Upstate Medical University), Dalila Pinto (Icahn School of Medicine at Mount Sinai), Panos Roussos (Icahn School of Medicine at Mount Sinai), Stephan Sanders (University of California, San Francisco), Nenad Sestan (Yale University), Pamela Sklar (Icahn School of Medicine at Mount Sinai), Matthew State (University of California, San Francisco), Patrick Sullivan (University of North Carolina), Flora Vaccarino (Yale University), Daniel Weinberger (Lieber Institute for Brain Development), Sherman Weissman (Yale University), Kevin White (University of Chicago), Jeremy Willsey (University of California, San Francisco), and Peter Zandi (Johns Hopkins University). We are grateful to Prof. Laurence A. Bindoff and Dr. Shreejoy Tripathy for the valuable comments and discussions.

## Contributions

L.T. conceived, designed and performed the ChIP-Seq and integrative computational analyses. G.T.T. designed and performed the immunoblots and cell-culture experiments. J.S. contributed to conception and implementation of the ChIP-seq and immunoblot analyses. K.H. contributed to data interpretation and critical revision of the manuscript. G.S.N. performed the preprocessing of the ChIP-seq dataset and contributed to the computational analyses. C.D. conceived, designed, supervised the immunoblots and cell-culture experiments, performed part of the immunoblot experiments, contributed to data analyses and interpretation. O.B.T. and G.A. contributed part of the tissue material, supplementary data, and critical revision of the manuscript. C.T. conceived, designed and directed the study, contributed to data analyses and interpretation and acquired funding for the study. L.T., G.T.T., C.D. and C.T. wrote the manuscript with the active input and participation of J.S. and G.S.N. All authors have read and approved the manuscript.

## Ethics declarations

### Ethics considerations

Informed consent was available from all individuals. Ethical permission for these studies was obtained from our regional ethics committee (REK 2017/2082, 2010/1700, 131.04)

