## Supplement methods, tables and figures for "Genome-wide dysregulation of histone acetylation in the Parkinson’s disease brain"

**TABLES**:

**Supplementary Table S1.** Cohort demographic information and experimental allocation.

**Supplementary Table S2.** Statistics of WB analyses

**Supplementary Table S3.** Peak number and genomic coverage in PD and controls

**Supplementary Table S4.** Differential ChIP-seq analysis results PW

**Supplementary Table S5.** Differential ChIP-seq analysis results NBB

**Supplementary Table S6.** Replicated Genes

**Supplementary Table S7.** Replicated DARs (full list of the common regions)

**Supplementary Table S8.** PD-implicated genes

**Supplementary Table S9.** Association of -log10(p) H3K27 hyperacetylation (one sided p-value) with the number of p300 and non-SIRT1 HDAC binding sites

**Supplementary Table S10.** Association of -log10(p) differential H3K27ac with being a PD-implicated gene and EGL

**Supplementary Table S11.** Enrichment analysis of the most decoupled genes and most negatively correlated genes

**FIGURES:**

**Supplementary Figure S1**. Distribution of genomic annotations and p-values among unique and common peaks

**Supplementary Figure S2.** IGV track of H3K27ac ChIP-Seq peaks annotated to candidate housekeeping genes.

**Supplementary Figure S3.** Correlation of the total number of reads from ChIP-seq normalized by different measures with Western blot data from the same subjects

**Supplementary Figure S4.** Validation of MSP approach for estimation of relative cellular abundance across samples

**Supplementary Figure S5**. Cellular composition estimated by MSPs in cortical samples of PD patients

**Supplementary Figure S6.** Significance of association of different variables with the first five principal components of the data

**Supplementary Figure S7.** Replication of ChIP-seq analysis in NBB cohort

**Supplementary Figure S8.** Decreased correlation between promoter H3K27ac state and gene expression in PD in RLE-normalized ChIP-seq data

**Supplementary methods**

**Chemicals, reagents and media**

Unless otherwise indicated, all chemicals were from Sigma Aldrich or from Bio-Rad. Cell culture media and reagents were from Invitrogen.

**Lysate preparation, protein determination and immunoblotting**

Fresh frozen brain tissue was lysed in 50 mM Tris-HCl pH 8.0, 150 mM NaCl, 1 mM EDTA and 2% SDS. SH-SY5Y cells were lysed in 50 mM Tris-HCl pH 8.0, 150 mM NaCl, 1 mM EDTA, 0.2% SDS, 10 µM trichostatin A, 10 µM nicotinamide and complete protease inhibitor cocktail. All samples were homogenized with a 27G syringe and centrifuged at 12.000 x g for 5 min at room temperature. Supernatants were collected and protein concentration determined using the BCA protein kit (Pierce/Thermofisher). Three independently dissected samples of brain tissue (replicates) were analyzed from each individual. Immunoblots were repeated at least 2 times for each marker.

For immunoblotting, 15 µg total protein were separated using AnykD^TM^ MiniProtean® TGX^TM^ Precast Protein gradient gels (Bio-Rad) and transferred to a 0.2 µm PVDF membrane using the Trans-Blot Turbo system (Bio-Rad) at 25 V and 1 Amp for 30 min. After transfer, membranes were blocked with 5% BSA in TBS containing 0.1 % Tween-20 (TBS-Tween) for antibodies targeting acetylated lysines and in 5% nonfat drymilk in TBS-Tween for all other antibodies. Incubation with primary antibody was carried out overnight at 4 ^o^C. Primary antibodies were used against acetylated lysine (CST #9441, 1:10.000), histone H2AK5ac (Abcam, ab1764, 1:2000), histone H2BK15ac (CST #9083, 1:10.000), histone H3K9K14ac (CST #9677, 1:20.000), histone H3K27ac (CST #8173S, 1:1000), histone H3K56ac (CST #4243, 1:200), histone H4K5ac (CST #8647, 1:1000), histone H4K12ac (CST #13944S, 1:2000), histone H4K16ac (CST #13534, 1:5000), histone H2A (Abcam, ab18255, 1:1000), histone H2B (Abcam, ab1790, 1:10.000), histone H3 (Abcam, ab1791m 1:20.000), histone H4 (CST # 2935, 1:500), GAPDH (SantaCruz, sc-32233, 1:20.000), NeuN (Millipore, MAB377, 1:1000), GFAP (Sigma-Aldrich, G3893, 1:15.000), CNP1 (Atlas antibodies, AMAb91072, 1:4000), CX3CR1 (Abcam, ab8021, 1:500), NUP188 (Invitrogen, PA5-48940, 1:250), calbindin (Sigma-Aldrich C9848, 1:3000), α-tubulinK40ac (Abcam, ab179484, 1:2000), α-tubulin (Abcam, ab18251, 1:2000), SIRT1 (Sigma Aldrich, S5447, 1:250), SIRT2 (Sigma Aldrich, S8447, 1:2000), SIRT3 (Sigma Aldrich, S4072, 1:2000).

After washing four times with TBS-Tween for 5 min, HRP-conjugated secondary antibodies (rabbit anti-mouse (P0260) or swine anti-rabbit (P0217), DAKO) were incubated for one hour at RT. After washing three times with TBS-Tween and once with TBS for 5 min, immunodetection was carried out using enhanced chemiluminescence (Clarity ECL detection kit, Bio-Rad), images were taken on a ChemiDoc XRS imager (Bio-Rad) and bands were quantified by densitometry using the ImageLab software (Bio-Rad).

To normalize the histone acetylation signal to histone quantity, each acetylation blot was stripped twice using stripping buffer containing 0.2 M glycine, 3.5 mM SDS and 1% Tween-20 at pH 2.2 for one hour at 50 ^o^C. Complete stripping was confirmed by incubation of the membranes with secondary antibodies only and ECL. No bands were detected even after long exposure time (up to 10 min). After detection, the membrane was washed and re-probed with primary antibody recognizing the histone being studied for assessment of total histone protein amount. To exclude potential bias due to incomplete stripping, we performed additional experiments where acetylated and total histone markers for each histone modification were assessed in parallel on different blots. To estimate the acetylated fraction of each histone marker, we calculated the signal ratio of acetylated marker to total histone protein. This ratio is referred to as the “acetylation fraction” for each histone marker.

**Cell culture**

SH-SY5Y cells (ATCC® CRL-2266^TM^) were cultivated in DMEM /F-12 (1:1) medium with GlutaMAX containing 10 % FBS and penicillin / streptomycin. For differentiation, 8 x 10^5^ cells were seeded in a 6-well plate, and at about 80 % confluency sequentially treated with 10 µM retinoid acid in complete medium for 5 days and serum-free medium containing 50 ng / ml BDNF (Tocris) for 3 days as described previously^1^. For pharmacological treatment, fully differentiated cells were exposed to commonly prescribed anti-Parkinson drugs (APDs) including L-dopa (50 µM), ropinirole (0.05 µM), entacapone (0.05 µM and 0.5 µM) and carbidopa (30 µM and 1.5 µM,) for 48 h, or deacetylase inhibitor trichostatin A (TSA, 0.5 µM) for 8 hours. The applied concentrations were calculated based on the maximum clinical dose according to the Norwegian prescription catalogue (<https://www.felleskatalogen.no/medisin>). The maximum prescribed dosage was converted to blood concentration, according to reported pharmacokinetic properties in the Norwegian prescription catalogue, and used in the experiment. When the calculated values were below the described IC50 of the compound, an additional sample using the IC50 concentration was included. For L-DOPA, the concentration was chosen based on previous reports^2^. Cell viability was assessed by resazurin assay to optimize drug exposure time. Each cell culture experiment was conducted three times starting with undifferentiated SH-SY5Y cells.

**Chromatin Immunoprecipitation sequencing (ChIP-seq)**

For each ChIP-seq experiment approximately 200mg of frozen brain tissue per sample was fixed in PBS + 1% formaldehyde at room temperature for 15 minutes. Fixation was stopped by the addition of glycine to final concentration 0.125 M. The tissue pieces were then treated with a TissueTearer and finally spun down and washed 2x in PBS. Tissue was then homogenized with a lysis buffer using a manual Dounce homogenizer. Lysates were sonicated and the DNA sheared to an average length of 300-500 bp. 14 ug of chromatin was precleared with protein A agarose beads (Invitrogen). Protein-DNA complexes were immuno-precipitated using 4 ug of H3K27Ac antibody (Active Motif, cat# 39133, lot# 01518010). Complexes were washed, eluted from the beads with SDS buffer, followed by de-crosslinking with proteinase K and RNase treatment overnight at 65 C. The ChIP DNA was purified by phenol-chloroform extraction and ethanol precipitation. Quantitative PCR (qPCR) reactions were carried out in triplicate on specific genomic regions using SYBR Green Supermix (Bio-Rad). The resulting signals were normalized for primer efficiency by carrying out qPCR for each primer pair using Input DNA. Illumina sequencing libraries were prepared from the ChIP and Input DNAs by the standard consecutive enzymatic steps of end-polishing, dA-addition, and adaptor ligation. After a final PCR amplification step, the resulting DNA libraries were quantified and sequenced on Illumina’s NextSeq 500 (75 base pair reads, single end). ChIP-seq was performed by Active Motif (<https://www.activemotif.com/>).

**Peak calling and read counting**

Peak calling procedure was carried out in two complementary approaches: 1) **Per group (PD or controls):** the peaks are called separately for each group, aggregating the relevant samples and inputs. While this approach is not optimal for differential peak calling analysis as it requires harmonization of peaks between the two groups, it can be used to compare the total genome coverage as well as unique H3K27ac genomic regions in each group. 2) **All samples combined:** the peaks are called by aggregating all samples and input controls. This approach does not allow identification of unique genomic regions however, it provides a better identification of consensus peaks, and thus, was used for differential peak analysis.

Specifically, BED files for single samples were merged into the three groups: (a) all PD samples, (b) all control samples (c) all samples. MACS2 version 2.1.1^68^ was used on each of the three group BED files without the shifting model (--nomodel) and with the following parameters: sequencing tag size (-s) 75, read extension (--extsize) 200, read shift (--shift) 0, and keeping all duplicates (--keep-dup all). The resulting peak-sets were filtered using the ENCODE black listed regions and only peaks within canonical chromosomes were kept for downstream analyses. Sample-specific enrichment on the identified peaks was performed using featureCounts version 1.6.4^69^ with default parameters. To this end, reads within filtered BAM files were extended to reach the fragment length (200) before quantification. For both the individual group and all samples combined approaches, peaks with non-adjusted p-value > 10^-7^ were excluded from the analysis. Reads inside peaks (RiP) were quantified using “featureCounts” program from the “subread” package v2.0.0.

**Calculation of Marker Site Profiles (MSP)**

In order to identify brain cell-type specific H3K27ac regions we first analyzed H3K27ac ChIP-seq data from NeuN^+^ and NeuN^-^ brain cells^25^. Cell-type based broadPeaks (CellType_peak-set), BAM files and metadata files were downloaded from <https://www.synapse.org/#!Synapse:syn4566010>. Differential regions between NeuN^+^ (neurons) and NeuN^-^ (glia) cells were calculated using “DESeq2” R package, including chromatin amount, library batch, sex, hemisphere, age and pH as covariates in the model. Peaks were defined as cell-type specific differentially acetylated regions (DARs) if they met the following criteria: 1) |fold change| > 4 and 2) mean count > 1000. Peaks were annotated to genes using the “build_annotations” function from “annotatr” R package based on UCSC hg19 genome assembly. Peaks were annotated to all genes for which they intersected a region between 5kb upstream from their transcription start site (TSS) to the end of their 5’UTR. In the next step, we intersected the genes with DARs between glia and neurons with expression-based cortical marker gene lists based on NeuroExpresso database^27^. The DARs were next reassigned to specific glial and neuronal cells if they were annotated to genes defined as cell-type specific based on NeuroExpresso. For example, all DARs annotated to *MBP*, defined as oligodendrocyte marker gene based on NeuroExpresso, were defined as oligodendrocyte marker sites (MSS). In the next step, the reads from our samples were quantified in regions defined by the CellType_peak-set. The corresponding RiP were then converted to counts per million (CPM) and transformed to log2(CPM + 1). Next, for each cell-type specific MSS, we performed principal component analysis based on the relevant peaks using “prcomp” function from the “stats” R package using (scale = T), as described in^27^. Marker Site Profiles (MSP) were defined as the scores of the samples in the first principal component, transformed to {0-1} range for visualization purposes.

**Identification of differentially acetylated regions**

In order to further increase the reliability of analyzed peaks, for differential peak analysis, we removed peaks with maximal coverage of < 5 reads/base, calculated as max(Peak_counts_)/(Peak_length_). Housekeeping normalization factor was calculated as follows: for each peak annotated to a house-keeping gene with mean count > 150 (housekeeping peak), for each sample, we first calculated the ratio between the sample count to the mean count of the peak across all samples (counts_sample_/counts_mean_). The housekeeping normalization factor was then defined as mean of the ratios for all relevant housekeeping peaks. Housekeeping normalization factors were calculated based on two sets of housekeeping genes: 1) housekeeping genes suggested based on stable transcript expression^24^ and 2) housekeeping genes manually selected based on known high and global expression level and very high and broad peak signal covering most of the gene body.

Differential peak analysis was performed using “nbinomWaldTest” function from “DESeq2” R package. Dispersions were calculated by using “estimateDispertions” function from “DESeq2” R package, by setting the fitType parameter to “local”. Normalization factors calculated by the “estimateSizeFactors” function were replaced with the housekeeping normalization factor based on our manually selected housekeeping gene set (*ACTB, GAPDH, UBC*). Peaks were annotated to genes as described in the previous section. The output of the analysis was obtained using DESeq2 “results” function, setting the cutoff for adjusted p-value to 0.05, and independentFiltering = T.

**RNA-sequencing**

Total RNA was extracted from prefrontal cortex tissue homogenate for all samples using RNeasy plus mini kit (Qiagen) with on-column DNase treatment according to manufacturer’s protocol. Final elution was made in 65 µl of dH2O. The concentration and integrity of the total RNA was estimated by Ribogreen assay (Thermo Fisher Scientiﬁc), and Fragment Analyzer (Advanced Analytical), respectively and 500 ng of total RNA was used for downstream RNA-seq applications. First, rRNA was removed using Ribo-Zero™ Gold (Epidemiology) kit (Illumina, San Diego, CA) using manufacturer’s recommended protocol. Immediately after the rRNA removal the RNA was fragmented and primed for the ﬁrst strand synthesis using the NEBNext First Strand synthesis module (New England BioLabs Inc., Ipswich, MA). Directional second strand synthesis was performed using NEBNExt Ultra Directional second strand synthesis kit. Following this the samples were taken into standard library preparation protocol using NEBNext® DNA Library Prep Master Mix Set for Illumina® with slight modiﬁcations. Brieﬂy, end-repair was done followed by poly(A) addition and custom adapter ligation. Post-ligated materials were individually barcoded with unique in-house Genomic Services Lab (GSL) primers and ampliﬁed through 12 cycles of PCR. Library quantity was assessed by Picogreen Assay (Thermo Fisher Scientiﬁc), and the library quality was estimated by utilizing a DNA High Sense chip on a Caliper Gx (Perkin Elmer). Accurate quantiﬁcation of the ﬁnal libraries for sequencing applications was determined using the qPCR-based KAPA Biosystems Library Quantiﬁcation kit (Kapa Biosystems, Inc.). Each library was diluted to a ﬁnal concentration of 12.5 nM and pooled equimolar prior to clustering. 125 bp Paired-End (PE) sequencing was performed on an Illumina HiSeq2500 sequencer (Illumina, Inc.). RNA quality, as measured by the RNA integrity number (RIN), varied across samples (mean = 5.3, range = 3.0-7.2 for PW; mean = 6.8, range = 3.2-9.1 for NBB, S5 File), although the difference between conditions did not reach statistical significance in any of the cohorts (t-test P = 0.72 and 0.90 for PW and NBB cohorts, respectively).

**RNA-seq analyses**

FASTQ files were trimmed using Trimmomatic version 0.36^3^ with the following parameters: ILLUMINACLIP:truseq.fa:2:30:10 LEADING:3 TRAILING:3 SLIDINGWINDOW:4:15. FASTQ ﬁles were assessed using fastQC version 0.11.5^4^ prior and following trimming. Transcript-level quantification was performed using Salmon version 0.9.1^5^ accounting for GC bias (--gcBias) and with inward-stranded-reverse library option (-l ISR) against the Ensembl release 75 transcriptome. Collapsing transcript-level quantiﬁcation to gene-level values was achieved using the tximport R package version 1.8.0^6^. Non-standard chromosomes and transcripts encoded by the mitochondrial genome were excluded from the analyses. In addition, transcripts were flagged for removal if they gathered more than 1% of the reads on more than half of the samples, which resulted in the removal of 3 and 4 transcripts from the PW and NBB cohorts, respectively. Similarly, low-expressed genes (i.e. genes whose expression was below the median expression in at least 20% of the samples) were filtered out from downstream analyses. Samples were then marked as outliers if their median correlation in gene expression (log counts per million) with the other samples was below Q1-1.5*IQR or above Q3+1.5*IQR (Tukey’s fences; Q1: first quartile, Q3: third quartile, IQR: inter-quartile range). As a result, 3 samples were marked as outliers in the PW cohort and 3 in the NBB cohort, and were not included in downstream analyses (resulting sample sizes: N = 26, N = 18, for PW and NBB cohorts, respectively).

**Supplementary Table S2. Statistics of WB analyses**

|  |  | Corrected for age, sex, PMI | | Corrected for age, sex, PMI and cell composition | |
| --- | --- | --- | --- | --- | --- |
| Region | **Marker** | **β** | **p-value** | **β** | **p-value** |
| Prefrontal cortex | **Lys-Ac:GAPDH** | **0.27** | **0.014** | **0.54** | **0.006** |
|  | α-tubulinK40c:α-tubulin | -0.04 | 0.713 | -0.17 | 0.172 |
|  | H2AK5ac:H2A | -0.15 | 0.160 | -0.16 | 0.203 |
|  | **H2BK15ac:H2B** | **0.37** | **1.7E-05** | **0.37** | **1.6E-04** |
|  | **H3K9/14ac:H3** | **0.41** | **3.9E-04** | **0.48** | **3.9E-04** |
|  | **H3K27ac:H3** | **0.40** | **4.5E-08** | **0.42** | **8.3E-07** |
|  | **H3K56ac:H3** | **0.22** | **0.010** | **0.31** | **0.002** |
|  | H4K5ac:H4 | -0.08 | 0.345 | -0.10 | 0.299 |
|  | **H4K12:H4** | **0.30** | **0.005** | **0.38** | **0.003** |
|  | H4K16ac:H4 | -0.14 | 0.202 | -0.08 | 0.492 |
|  | **H2A:GAPDH** | **0.37** | **2.7E-05** | **0.16** | **1.2E-05** |
|  | **H2B:GAPDH** | **0.41** | **1.1E-08** | **0.48** | **6.4E-09** |
|  | **H3:GAPDH** | **0.30** | **2.0E-07** | **0.37** | **8.4E-08** |
|  | **H4:GAPDH** | **0.38** | **1.9E-18** | **0.44** | **1.5E-18** |
|  | **SIRT1:β-tubulin** | **0.30** | **0.002** | **0.35** | **0.002** |
|  | SIRT2:β-tubulin | 0.05 | 0.567 | -0.23 | **0.024** |
|  | **SIRT3:β-tubulin** | **0.27** | **0.003** | **0.35** | **0.001** |
|  | NUP188:GAPDH | 0.15 | 0.163 |  |  |
|  | NeuN:GAPDH | -0.14 | 0.050 |  |  |
|  | GFAP:GAPDH | 0.19 | 0.062 |  |  |
|  | **CNP1:GAPDH** | **0.38** | **3.9E-05** |  |  |
|  | CX3CR1:GAPDH | 0.00 | 0.994 |  |  |
| Striatum | H3K9/14ac:H3 | -0.06 | 0.695 | -0.02 | 0.912 |
|  | **H3K27ac:H3** | **0.20** | **0.034** | 0.11 | 0.258 |
|  | **H3:GAPDH** | **0.30** | **7.9E-05** | **0.25** | **0.002** |
|  | NeuN:GAPDH | -0.12 | 0.227 |  |  |
|  | GFAP:GAPDH | 0.18 | 0.135 |  |  |
|  | CNP1:GAPDH | 0.16 | 0.146 |  |  |
|  | **CX3CR1:GAPDH** | **-0.30** | **0.013** |  |  |
| Cerebellum | **H3K9/14ac:H3** | **-0.33** | **0.007** | **-0.29** | **0.013** |
|  | **H3K27ac:H3** | **0.20** | **0.016** | **0.25** | **0.005** |
|  | H3:GAPDH | 0.02 | 0.802 | 0.00 | 0.973 |
|  | **Calbindin:GAPDH** | **0.27** | **0.048** |  |  |
|  | GFAP:GAPDH | 0.16 | 0.096 |  |  |
|  | CNP1:GAPDH | -0.25 | 0.057 |  |  |
|  | CX3CR1:GAPDH | 0.05 | 0.698 |  |  |

Significant differences between individuals with PD and controls are highlighted.

**Supplementary Table S3. Peak number and genomic coverage in PD and controls**

|  | Condition | Total Peaks | Total Coverage | Common Peaks | Unique Peaks | Unique Peaks (%) | Total Coverage (%) |
| --- | --- | --- | --- | --- | --- | --- | --- |
| PW | PD | 146,763 | 2.85E+08 | 99,434 | 47,329 | 32.2 | 10.6 |
|  | Controls | 135,662 | 2.46E+08 | 117,822 | 17,840 | 13.2 | 9.11 |
| NBB | PD | 120,726 | 2.35E+08 | 93,229 | 27,497 | 22.8 | 8.7 |
|  | Controls | 117,800 | 2.32E+08 | 93,312 | 24,488 | 20.8 | 8.59 |

*Common peaks are peaks overlapping by at least one base between the groups

**Supplementary tables S4-S7 and S11 are available online and through -** [**https://github.com/ltoker/ChIPseqPD**](https://github.com/ltoker/ChIPseqPD)

**Supplementary Table S8. Altered H3K27 acetylation of PD implicated genes**

| GeneSymbol | GWAS  Nalls2019 | Monogenic PD | PD neuropathology | Adj  MetaP | AdjPval  PW | AdjPval  NBB |
| --- | --- | --- | --- | --- | --- | --- |
| *DLG2* | YES |  |  | **3.70E-07** | **9.80E-04** | **2.00E-03** |
| *MAP4K4* | YES |  |  | **1.30E-04** | **0.02** | **5.30E-03** |
| *CRHR1* | YES |  |  | **1.80E-04** | **0.04** | **3.30E-03** |
| *APP* |  |  | YES | **5.30E-04** | **0.02** | **0.01** |
| *MBNL2* | YES |  |  | **6.40E-04** | **9.50E-03** | **7.20E-03** |
| *STK39* | YES |  |  | **2.30E-03** | 0.18 | **2.50E-03** |
| *SH3GL2* | YES |  |  | **5.60E-03** | **0.03** | **0.05** |
| *MAPT* |  |  | YES | **9.20E-03** | **0.03** | 0.23 |
| *FYN* | YES |  |  | **0.01** | 0.22 | **8.70E-03** |
| *GPNMB* | YES |  |  | **0.01** | 0.23 | **0.03** |
| *NUCKS1* | YES |  |  | **0.01** | **0.03** | 0.33 |
| *SPPL2B* | YES |  |  | **0.01** | **0.02** | 0.08 |
| *DYRK1A* | YES |  |  | **0.02** | **0.03** | 0.46 |
| *GS1-124K5.11* | YES |  |  | **0.02** | 0.11 | 0.06 |
| *SNCA* | YES | YES | YES | **0.02** | 0.15 | 0.07 |
| *PARK7* |  | YES |  | **0.02** | **0.03** | 0.27 |
| *CAB39L* | YES |  |  | **0.03** | 0.13 | 0.14 |
| *MCCC1* | YES |  |  | **0.03** | 0.09 | 0.12 |
| *PRKN* |  | YES |  | **0.03** | 0.16 | **0.05** |
| *MED12L* | YES |  |  | **0.04** | 0.26 | **0.03** |
| *RNF141* | YES |  |  | **0.04** | 0.06 | 0.29 |
| *FAM171A2* | YES |  |  | **0.05** | 0.12 | 0.14 |
| *FBRSL1* | YES |  |  | **0.05** | 0.09 | **0.05** |
| *FCGR2A* | YES |  |  | **0.05** | 0.46 | **0.05** |
| *TMEM175* | YES |  |  | **0.05** | 0.11 | 0.24 |
| *RPS6KL1* | YES |  |  | 0.06 | 0.1 | 0.16 |
| *CHRNB1* | YES |  |  | 0.07 | 0.07 | 0.13 |
| *KCNIP3* | YES |  |  | 0.07 | 0.12 | 0.16 |
| *PINK1* |  | YES |  | 0.07 | 0.14 | 0.27 |
| *FAM49B* | YES |  |  | 0.08 | 0.21 | 0.19 |
| *SETD1A* | YES |  |  | 0.08 | 0.06 | 0.39 |
| *SYT17* | YES |  |  | 0.08 | 0.09 | 0.21 |
| *SATB1* | YES |  |  | 0.09 | 0.16 | 0.23 |
| *GAK* | YES |  |  | 0.1 | 0.07 | 0.29 |
| *GALC* | YES |  |  | 0.1 | 0.13 | 0.1 |
| *GBAP1* | YES |  |  | 0.1 | 0.08 | 0.75 |
| *IP6K2* | YES |  |  | 0.11 | 0.08 | 0.68 |
| *ASXL3* | YES |  |  | 0.12 | 0.19 | 0.33 |
| *SIPA1L2* | YES |  |  | 0.12 | 0.15 | 0.11 |
| *BST1* | YES |  |  | 0.14 | 0.1 | 0.86 |
| *CAMK2D* | YES |  |  | 0.14 | 0.34 | 0.11 |
| *BIN3* | YES |  |  | 0.16 | 0.25 | 0.25 |
| *GBF1* | YES |  |  | 0.16 | 0.1 | 0.43 |
| *CHD9* | YES |  |  | 0.17 | 0.08 | 0.25 |
| *CLCN3* | YES |  |  | 0.17 | 0.25 | 0.37 |
| *CTSB* | YES |  |  | 0.17 | 0.56 | 0.17 |
| *SCARB2* | YES |  |  | 0.18 | 0.18 | 0.39 |
| *WNT3* | YES |  |  | 0.2 | 0.12 | 0.5 |
| *DNAH17* | YES |  |  | 0.22 | 0.26 | 0.42 |
| *MEX3C* | YES |  |  | 0.23 | 0.61 | 0.2 |
| *FAM47E* | YES |  |  | 0.24 | 0.26 | 0.52 |
| *IGSF9B* | YES |  |  | 0.25 | 0.14 | 0.74 |
| *TMEM163* | YES |  |  | 0.25 | 0.28 | 0.49 |
| *UBAP2* | YES |  |  | 0.25 | 0.34 | 0.42 |
| *RPS12* | YES |  |  | 0.26 | 0.57 | 0.27 |
| *ITPKB* | YES |  |  | 0.27 | 0.22 | 0.69 |
| *INPP5F* | YES |  |  | 0.28 | 0.57 | 0.25 |
| *LCORL* | YES |  |  | 0.28 | 0.56 | 0.3 |
| *RIMS1* | YES |  |  | 0.28 | 0.27 | 0.59 |
| *ELOVL7* | YES |  |  | 0.3 | 0.62 | 0.3 |
| *PAM* | YES |  |  | 0.35 | 0.41 | 0.5 |
| *HIP1R* | YES |  |  | 0.37 | 0.29 | 0.66 |
| *SPTSSB* | YES |  |  | 0.38 | 0.97 | 0.26 |
| *KCNS3* | YES |  |  | 0.39 | 0.44 | 0.52 |
| *PMVK* | YES |  |  | 0.39 | 0.64 | 0.38 |
| *LRRK2* | YES | YES |  | 0.4 | 0.98 | 0.28 |
| *BAG3* | YES |  |  | 0.41 | 0.63 | 0.41 |
| *VAMP4* | YES |  |  | 0.42 | 0.36 | 0.68 |
| *SCAF11* | YES |  |  | 0.46 | 0.18 | 0.37 |
| *FAM47E-STBD1* | YES |  |  | 0.47 | 0.4 | 0.55 |
| *GCH1* | YES |  |  | 0.47 | 0.31 | 0.87 |
| *C5orf24* | YES |  |  | 0.55 | 0.05 | 0.47 |
| *RETREG3* | YES |  |  | 0.57 | 0.24 | 0.73 |
| *NOD2* | YES |  |  | 0.63 | 0.42 | 0.75 |
| *MIPOL1* | YES |  |  | 0.66 | 0.41 | 0.99 |
| *UBTF* | YES |  |  | 0.66 | 0.76 | 0.61 |
| *FGF20* | YES |  |  | 0.69 | 0.74 | 0.66 |
| *KRTCAP2* | YES |  |  | 0.77 | 0.5 | 0.99 |
| *VPS35* |  | YES |  | 0.8 | 0.99 | 0.64 |
| *RAB29* | YES |  |  | 0.82 | 0.79 | 0.79 |
| *CRLS1* | YES |  |  | 0.9 | 0.74 | 0.95 |
| *VPS13C* | YES |  |  | 0.9 | 0.87 | 0.84 |
| *KPNA1* | YES |  |  | 0.99 | 0.98 | 0.95 |
| *BRIP1* | YES |  |  | ---- | 0.23 | ---- |
| *CD19* | YES |  |  | ---- | 0.28 | ---- |
| *LINC00693* | YES |  |  | ---- | ---- | 0.98 |
| *HLA-DRB5* | YES |  |  | ---- | ---- | ---- |
| *ITGA8* | YES |  |  | ---- | ---- | ---- |
| *LOC100131289* | YES |  |  | ---- | ---- | ---- |
| *RIT2* | YES |  |  | ---- | ---- | ---- |
| *TRIM40* | YES |  |  | ---- | ---- | ---- |
| *CASC16* | YES |  |  | ---- | ---- | ---- |

Out of the 92 PD implicated genes, 83 were could be assessed by ChIP-seq data in both cohorts. In bold, adjusted metaP/p-value < 0.05.

--- indicates that differential H3K27 acetylation for the gene could not be assessed either due to lack of representation of the gene in ChIP-seq data or due to DESeq2 independent filtering.

**Supplementary Table S9.** Association of -log10(p) H3K27 hyperacetylation (one sided p-value) with the number of p300 and non-SIRT1 HDAC binding sites

|  |  | PW | | | NBB | | |
| --- | --- | --- | --- | --- | --- | --- | --- |
|  |  | Estimate | t value | pvalue | Estimate | t value | pvalue |
| Gene level | (Intercept) | 1.13 | 988.95 | 0.00E+00 | 0.58 | 542.56 | 0.00E+00 |
|  | EP300 | 0.01 | 40.08 | 0.00E+00 | 0.03 | 154.28 | 0.00E+00 |
|  | HDAC_Binding | -0.01 | -40.14 | 0.00E+00 | -0.04 | -154.17 | 0.00E+00 |
|  | PDgeneYes | 0.11 | 8.17 | 3.09E-16 | 0.09 | 7.43 | 1.12E-13 |
| Peak level | (Intercept) | 1.09 | 507.48 | 0.00E+00 | 0.78 | 321.24 | 0.00E+00 |
|  | EP300 | 0.02 | 7.90 | 2.71E-15 | 0.09 | 35.99 | 3.03E-282 |
|  | HDAC_Binding | -0.07 | -29.44 | 5.34E-190 | -0.14 | -56.52 | 0.00E+00 |
|  | PDgeneYes | 0.33 | 13.40 | 6.67E-41 | 0.16 | 5.90 | 3.75E-09 |

Output of linear model. HDAC_binding – the sum of bindings sites for HDAC1, HDAC2, HDAC6, HDAC8 and SIRT6. For the gene level analysis, binding sites were first quantified for each peak and then collapsed to genes. PDgene – PD_implicated gene.

**Supplementary Table S10. Association of -log(p) differential H3K27ac with being a PD implicated gene and EGL**

|  | -log10(metaP) | | -log10(pValue) PW | | -log10(pValue) NBB | |
| --- | --- | --- | --- | --- | --- | --- |
|  | PDgene | log10(EGL) | PDgene | log10(EGL) | PDgene | log10(EGL) |
| Hypoacetylated regions | β = -0.61,  p = 2e-39 | β = -0.24,  p = 2e-26 | β = 0.06,  p < 2e-05 | β = -0.01,  p = 2e-04 | β = -0.19,  p = 1e-26 | β = -0.22,  p < 2e-26 |
| Hyperacetylated regions | β = 0.16,  p = 1e-07 | β = 0.42,  p < 2e-26 | β = 0.1,  p = 4e-14 | β = 0.01,  p = 1e-05 | β = 0.11,  p = 4e-11 | β = 0.23,  p < 2e-26 |

Output of linear model adjusting for the number of peaks/gene. The results are based on common regions (evaluated for -log10(metaP)) or each cohort separately (evaluated for -log10(pValue)). PDgene – PD_implicated gene. EGL – effective gene length.

**Supplementary Figures**

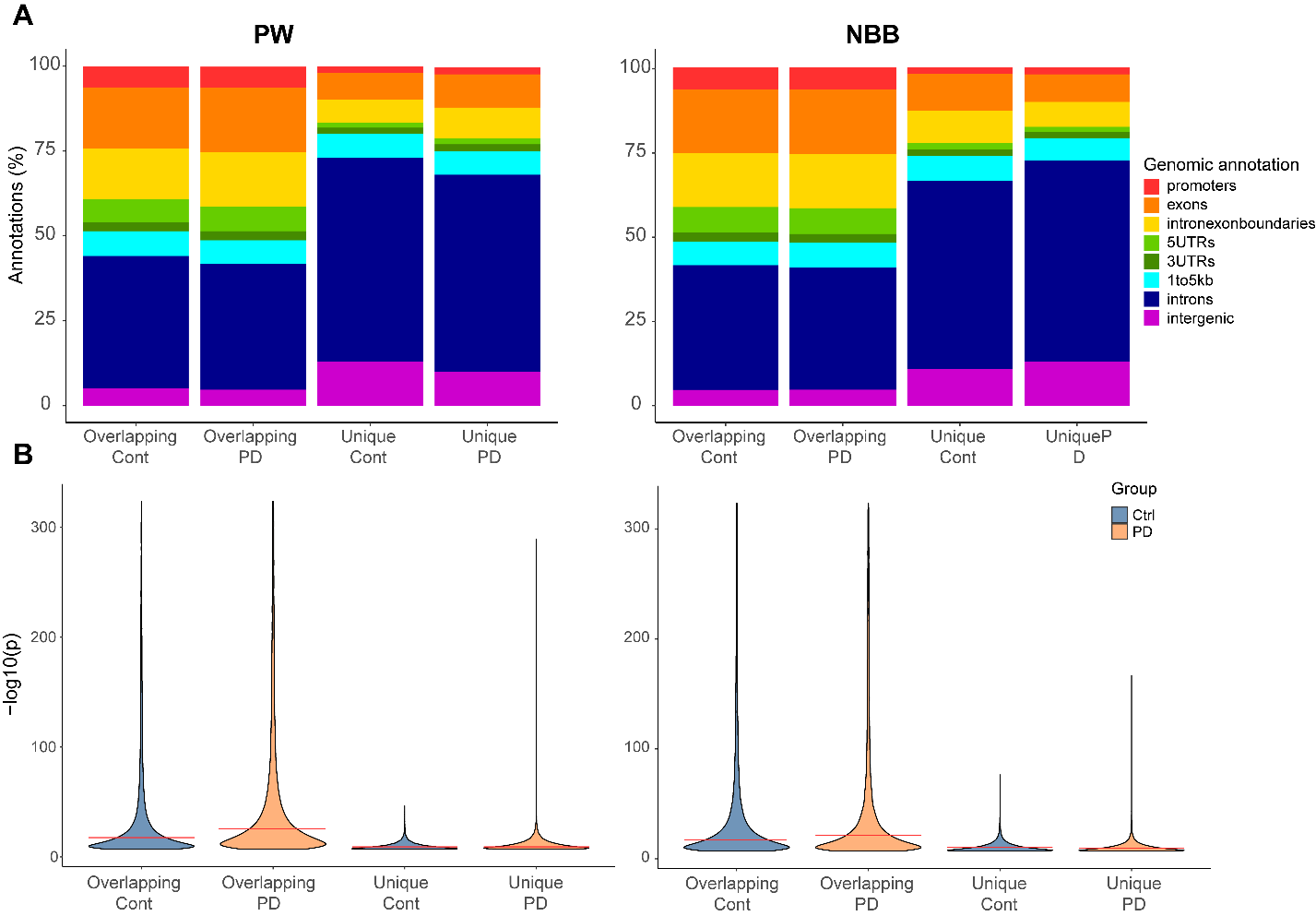

**Supplementary Figure S1. Distribution of genomic annotations and p-values among unique and common peaks**

**A.** Overlapping peaks (peaks with genomic overlap of at least one base) identified in both PD and control groups and unique peaks (peaks with no genomic overlap) were annotated to functional genomic region using the ‘annotatr’ R package. The percentage of annotations falling in each of the categories is shown on the y-axis. Results are shown for PW (left) and NBB (right) cohorts. 1to5kb: Region 5kb-1kb upstream to transcription start site (TSS).

**B.** Distribution of -log10(p-values) reported from MACS2 algorithm, compared to input control of unique and common peaks in PW (left) and NBB (right) cohorts. The median in each group is indicated by the red line. In both cohorts, unique peaks in either of the groups are characterized by higher p-values (wilcoxon’s p < 10^-16^). Overlapping Cont: common peaks in controls, Overlapping PD: overlapping peaks in PD, Unique Cont: unique peaks in controls, Unique PD: unique peaks in PD.

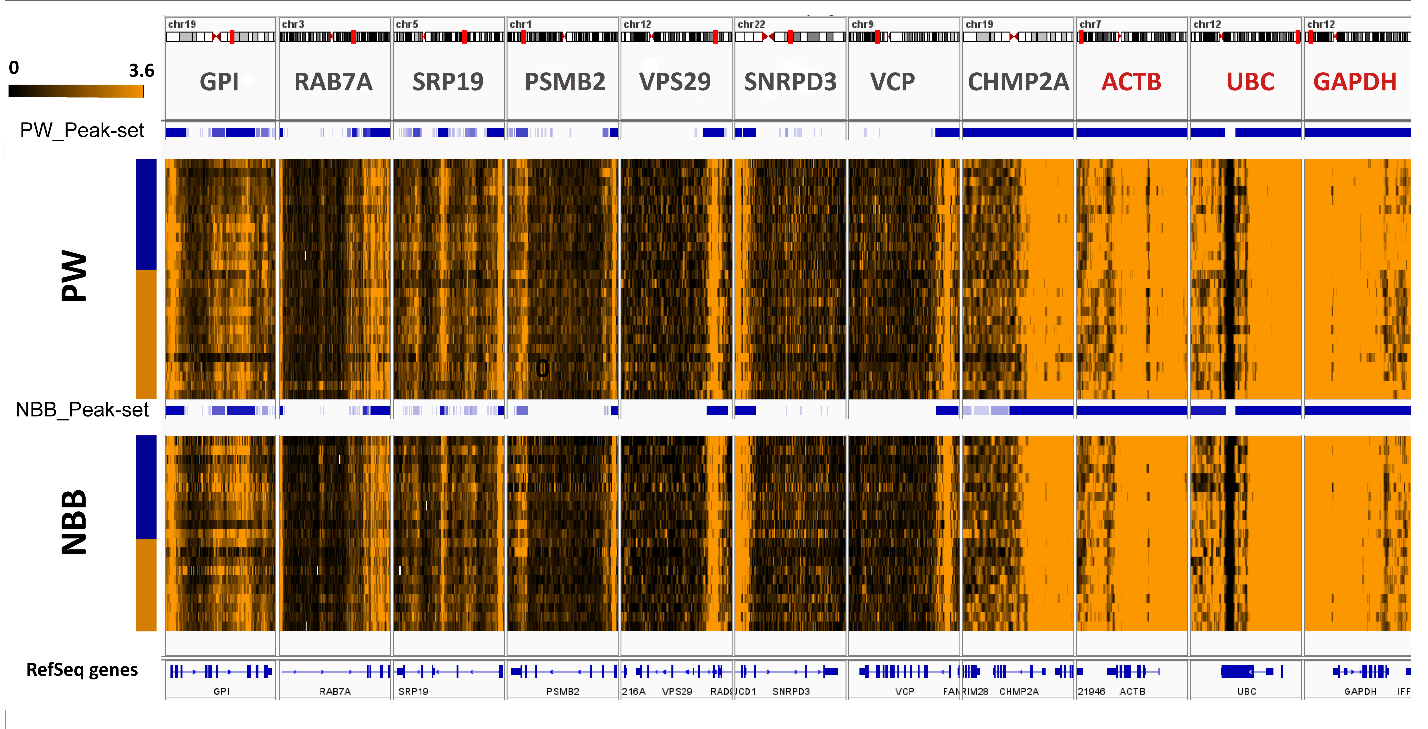

**Supplementary Figure S2. H3K27ac ChIP-Seq peaks annotated to candidate housekeeping genes.**

IGV representation of H3K27ac binding to genomic regions annotated to house keeping genes. Genes indicated in grey were suggested by Allhoff et al.^9^. Genes indicated in red were manually selected for the purpose of this study. H3K27ac peaks identified based on each cohort are indicated by blue rectangles. Blue horizontal boxes indicate regions identified by MACS2 algorithm as H2K27ac peaks based on PW or NBB peak-sets. The heatmaps are a representation of BigWig files for each sample (shown in rows), indicating the fold of change in coverage of each position, compared to input control. Blue and orange vertical side bars indicate control (blue) or PD (orange) samples. The RefSeq panel shows a graphical representation of the gene sequence.

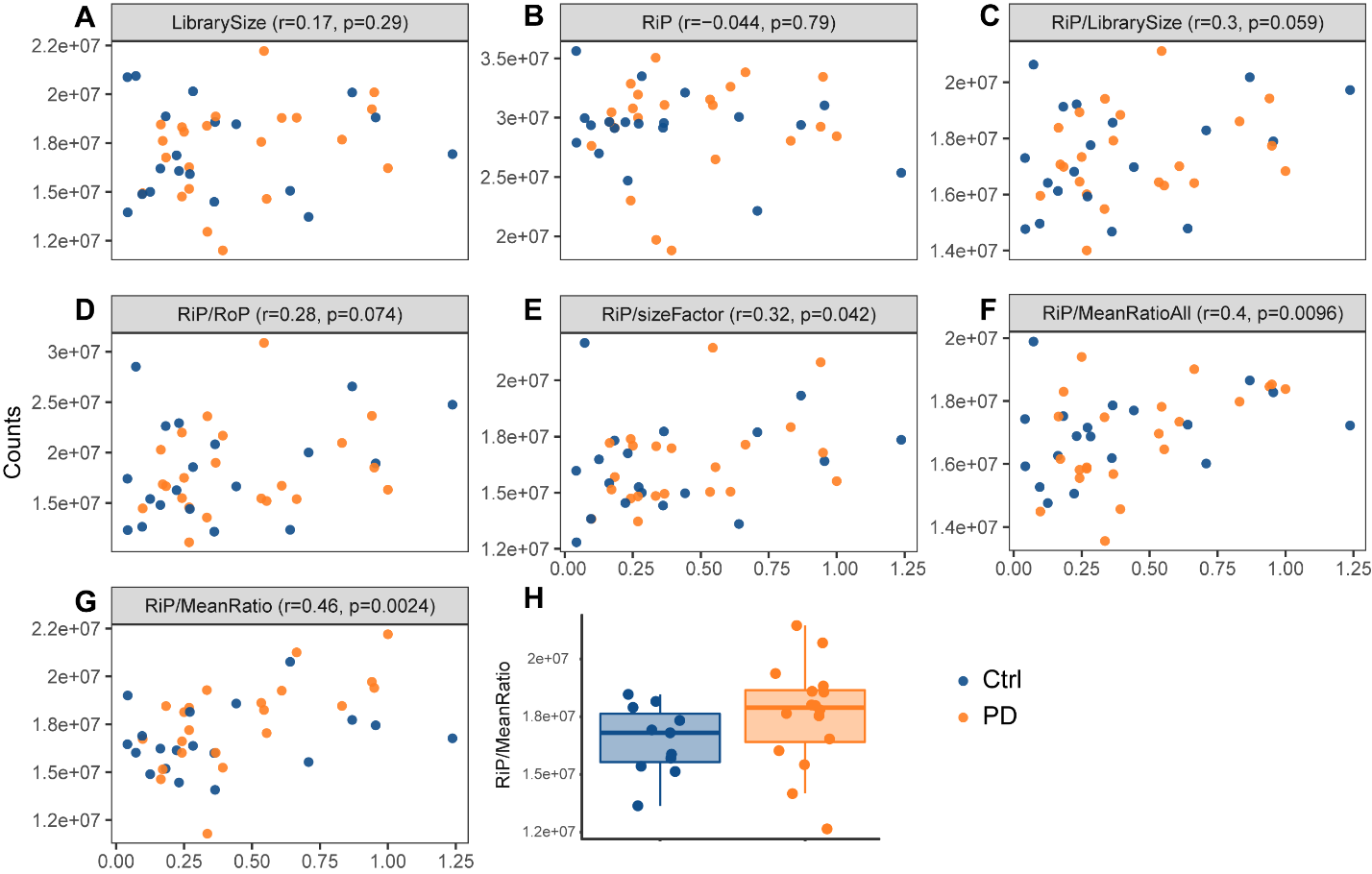

**Supplementary Figure S3. Correlation of the total number of reads from ChIP-seq data, normalized by different measures, with Western blot data from the same subjects.**

**A,B.** Correlation between the library size (**A**) or reads in peaks (**B**) and relative H3K27 acetylation levels normalized to GAPDH as determined by immunoblot analysis. **C-G,** Correlation between the total number of reads in peaks normalized by different measures and relative H3K27 acetylation levels normalized to GAPDH as determined by Western blot analysis. **H.** Group comparison of the total number of RiP normalized to house-keeping genes shows increase in total H3K27ac in PD.

LibrarySize – total number of reads; RiP – total number of reads in peaks; RoP – total number of reads outside the peaks (Librarysize - RiP); sizeFactor – RLE normalization factor, calculated by DESeq2; MeanRatioAll – geometric mean using peaks in genes suggested by Allhoff et al.^9^; MeanRatio – geometric mean of peaks annotated to our chosen housekeeping genes.

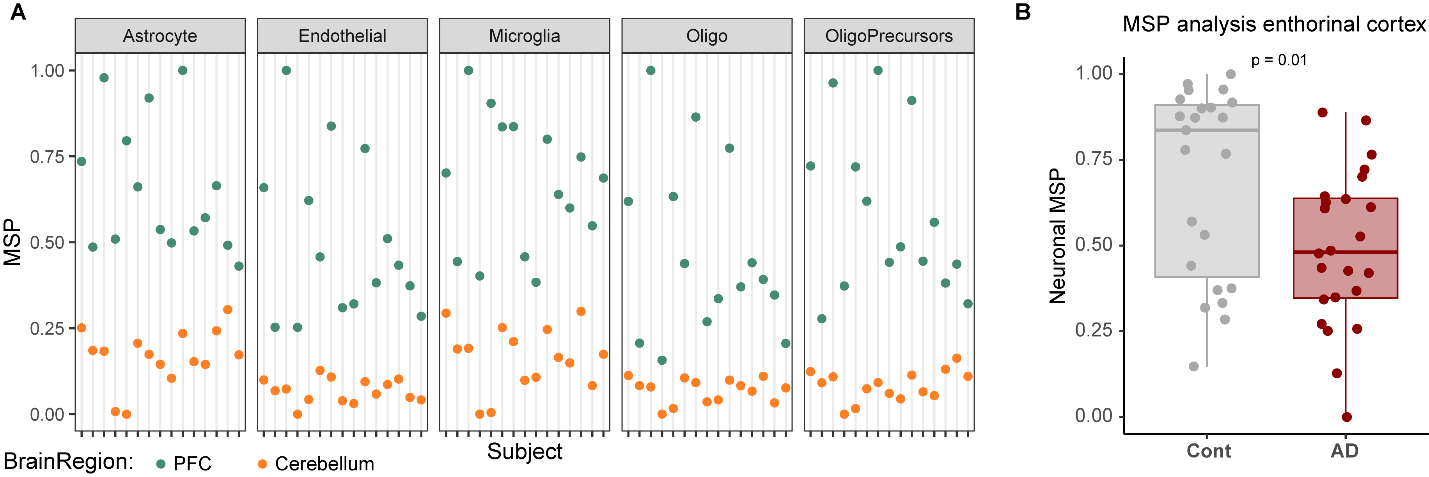

**Supplementary Figure S4. Validation of the MSP approach for estimation of relative cellular abundance across samples.**

**A.** Estimated MSPs in H3K27ac ChIP-seq data from prefrontal cortex (PFC, shown in green) or cerebellum (shown in orange) samples from 17 subjects, generated by Sun et al.^10^ . Oligo: oligodendrocytes. **B.** Estimated neuronal MSP in entorhinal cortex of subjects with Alzheimer’s disease (AD) and controls (Cont) generated by Marzi et al.^11^ . Group difference was assessed based on linear model adjusting for age and sex of the individuals.

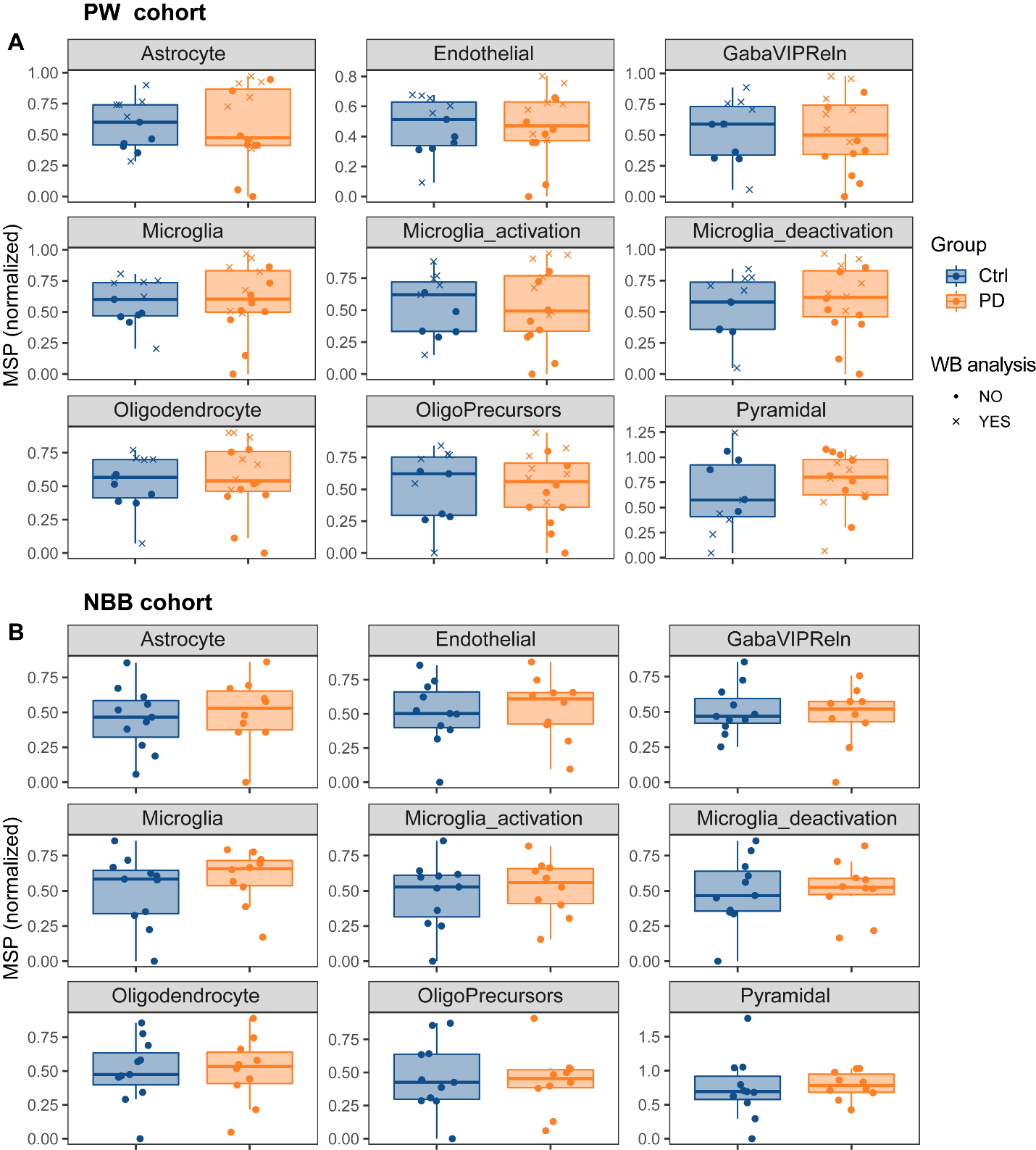

**Supplementary Figure S5. Cellular composition estimated by MSPs in cortical samples of PD patients.**

The estimated MSPs (normalized by the housekeeping peak ratio) are shown for PW (**A**) and NBB (**B**) cohorts. The protein levels of specific cell type markers were measured in a subset of individuals from the PW cohort (indicated by crosses). GabaViPReln: VIP and Reelin positive cells; Microglia_activation: genes upregulated in activated microglia; Microglia_deactivation: genes downregulated in activated microglia; OligoPrecursors: oligodendrocyte precursor cells

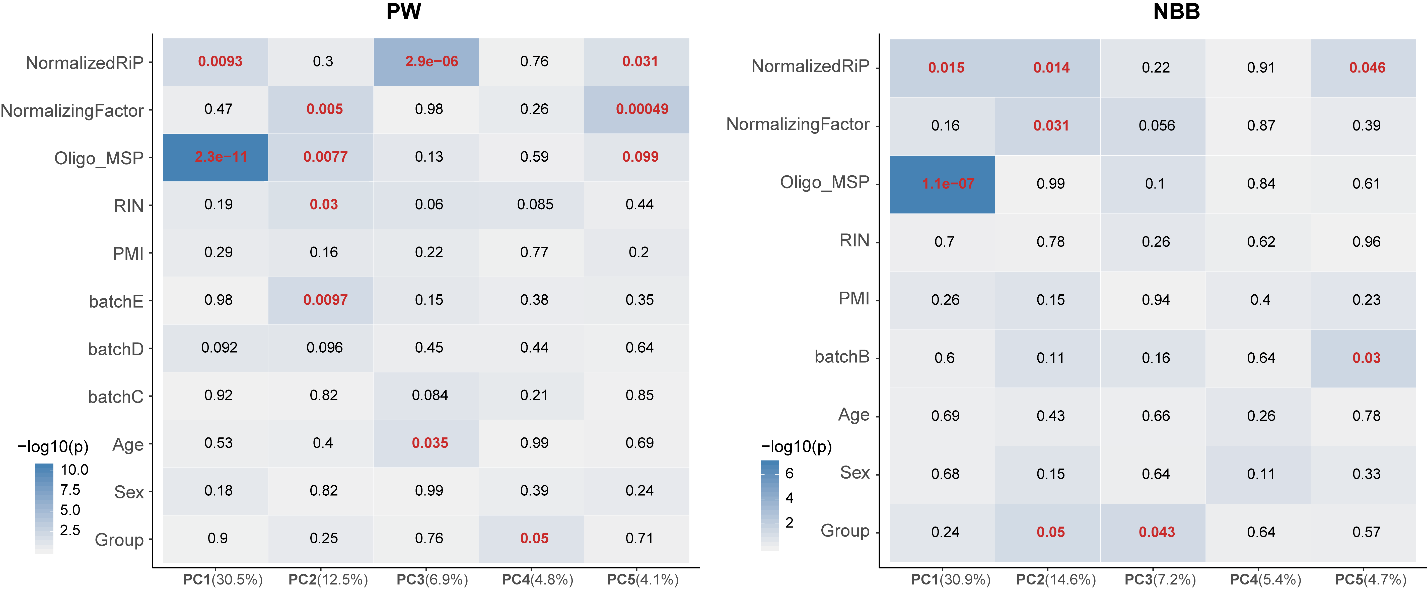

**Supplementary Figure S6. Significance of association of different variables with the first five principal components of the data.**

The strongest association in both cohort was found with the cellular composition of the samples, estimated by MSPs. The indicated numbers show the p-values of beta coefficients of the variables (indicated on the y-axis) for the first five PCs (indicated on the x-axis). Heatmap colors represent the -log10 of the p-values. Variance explained by each component is indicated in parentheses. Normalized RiP – effetive library size (total number of reads in peaks) normalized by the mean ratio of housekeeping peaks (normalizing factor).

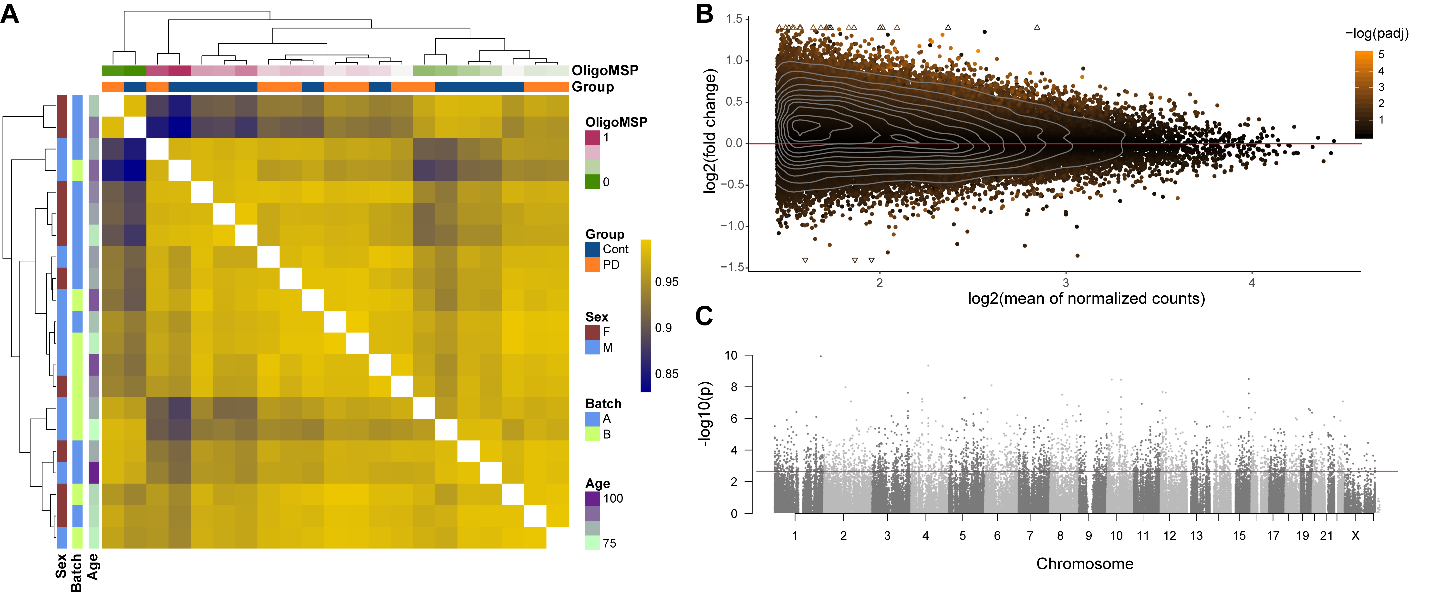

**Suplementary Figure S7. Replication of ChIP-seq analysis in the NBB cohort.**

**A.** Hierarchical clustering of the samples from the NBB cohort based on sample-to-sample correlation shows that, similar to the PW cohort, the samples cluster based on their cellular composition. Supplementary Figure S6 shows the association of each of the variables with the main principal components of the data. **B**. MA plot based on NBB_peak-set indicates that increased H3K27ac in PD is observed genome-wide, rather that being restricted to specific regions. **c**. Manhattan plot showing the distribution of genomic locations and differential p-values of the H3K27ac NBB_peak-set. The red dashed line indicates the -log10 of the highest p-value with < 5% false positive rate (Benjamini-Hochberg adjusted p-value < 0.05).

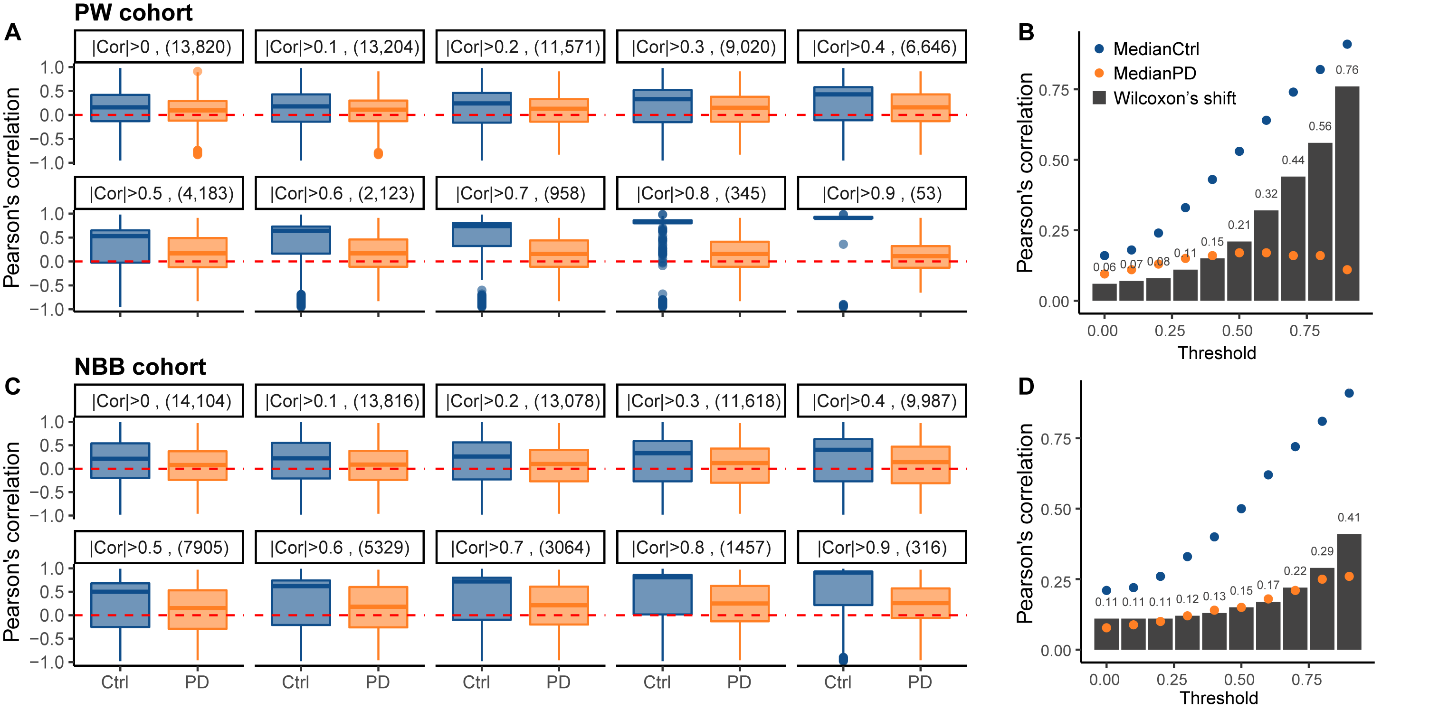

**Supplementary Figure S8. Decreased correlation between promoter H3K27 acetylation state and gene expression in PD in RLE-normalized ChIP-seq data.**

Pearson’s correlation between the adjusted promoter H3K27ac counts based on RLE-normalized ChIP-seq data and adjusted RNA-seq counts was calculated for each gene across control or PD individuals. The distribution of correlation was compared between the groups for various thresholds of minimal absolute correlation in either of the groups. Results are shown for PW (**A,B**) and NBB (**C,D**) cohorts. The number of genes that remained after applying the correlation threshold is shown in parentheses. The correlations remained closely distributed around 0 in PD subjects from both cohorts, regardless of the correlation threshold. **B,D** Median correlations in each group and the calculated Wilcoxon’s delta shift for different correlation thresholds.
